# Implantable nanophotonic neural probes for integrated patterned photostimulation and electrophysiology recording

**DOI:** 10.1101/2023.11.14.567101

**Authors:** Fu Der Chen, Homeira Moradi Chameh, Mandana Movahed, Hannes Wahn, Xin Mu, Peisheng Ding, Tianyuan Xue, John N. Straguzzi, David A. Roszko, Ankita Sharma, Alperen Govdeli, Youngho Jung, Hongyao Chua, Xianshu Luo, Patrick G. Q. Lo, Taufik A. Valiante, Wesley D. Sacher, Joyce K. S. Poon

## Abstract

Optogenetics has transformed neuroscience by allowing precise manipulation of neural circuits with light [1–5]. However, a central difficulty has been to deliver spatially shaped light and record deep within the brain without causing damage or significant heating. Current approaches form the light beam in free space and record the neural activity using fluorescence imaging or separately inserted electrodes [6–9], but attenuation limits optical penetration to around 1 mm of the brain surface [10]. Here, we overcome this challenge with foundry-fabricated implantable silicon neural probes that combine microelectrodes for electrophysiology recordings with nanophotonic circuits that emit light with engineered beam profiles and minimal thermal impact. Our experiments reveal that planar light sheets, emitted by our neural probes, excited more neurons and induced greater firing rate fatigue in layers V and VI of the motor and somatosensory cortex of Thy1-ChR2 mice at lower output intensities than low divergence beams. In the hippocampus of an epilepsy mouse model, we induced seizures, a network-wide response, with light sheets without exceeding the *∼* 1*^◦^*C limit for thermally induced electrophysiological responses [11–13]. These findings show that optical spatial profiles can be tailored for optogenetic stimulation paradigms and that the probes can photostimulate and record neural activity at single or population levels while minimizing thermal damage to brain tissue. The neural probes, made in a commercial silicon photonics foundry on 200-mm silicon wafers, demonstrate the manufacturability of the technology. The prospect of monolithically integrating additional well-established silicon photonics devices, such as wavelength and polarization multiplexers, temperature sensors, and optical power monitors, into the probes holds the potential of realizing more versatile, implantable tools for multimodal brain activity mapping.

## I. INTRODUCTION

Genetically encoded optogenetic actuators, inhibitors, and fluorescence indicators are essential tools in neuroscience that enable highly precise ways to dissect neural circuits with cell-type specificity, deterministic neuronal control, cellular spatial resolution, and millisecond time resolution [1–5]. These optical methods of neural activity mapping have motivated the generation of a wide variety of shaped light beams, such as planar sheets, focused spots, short pulses, and non-diffractive Bessel beams in the visible and near-infrared spectral range for delivery into brain tissue [6–9]. Typically, such light beams are formed in free space with discrete optical components, and their penetration depth into the brain is limited by tissue attenuation to about 100 *µ*m to 1 mm for wavelengths between 450 and 1100 nm [10]. Bringing light to deep brain regions beyond the attenuation length of light in mammalian brain tissues requires implantable light sources, such as optical fibers [14–16], miniature gradient index lenses [17–20] and silicon (Si) probes with optoelectronic devices (i.e., µ-LEDs or µ-OLED) [21–25] or waveguides with grating emitters [26–31].

The integration of microelectrodes and microscale light emitters on Si probes offers exciting experimental possibilities by enabling flexible and selective brain stimulation with light while recording the evoked electrophysiological responses [21, 24, 27, 28, 32, 33]. For probes that use waveguides, Libbrecht et al. realized foundry-fabricated neural probes with 12 optical outputs and 24 electrodes [27], but the need to mount laser diode carriers manually, one per optical channel, limited its practicality [27]. Mohanty et al. developed a Si photonic neural probe that was 250 *µ*m thick and several millimeters wide, incorporating a limited number of 4 electrodes and 8 light emitters [28], but the design was substantially larger than the accepted dimensions for minimizing brain tissue displacement (typically 70 to 100 *µ*m wide and *≤* 30 *µ*m thick) [21, 26, 27, 34, 35]. The most extensive integration to date employs µ-LEDs, with Si probes featuring up to 128 monolithically integrated µ-LEDs and 256 microelectrodes [24]. These probes offer a high density of light emitters unattainable by placing fibers or LEDs near electrophysiology probes [16, 36–38]. However, the low efficiency of the µ-LEDs (*<* 3% of the electrical power is converted to optical power) necessitates strict thermal management during operation [21, 22, 24, 39] and may prohibit these probes from being used in experiments requiring high optical powers for broad-area stimulation. Moreover, achieving optical beam profiles beyond the Lambertian emission of µ-LED probes for the control of stimulation volumes presents a significant challenge, necessitating the integration of additional micro-optic components such as lenses.

Here, we close the gap between tailored light profiles for photostimulation and moderate density electrophysiology recordings with implantable neural probes that monolithically integrate microelectrodes with photonic integrated circuits (PICs). The optical beam emission is generated by nanophotonic waveguide gratings on the probe, and different stimulation volumes can be targeted by a variety of optical profiles (e.g., planar sheets, low-divergence beams). In this study, the probes had either 18 electrodes and 16 low-divergence emitters on 1 shank, or 56 electrodes and 5 light sheet emitters on 4 shanks. We have previously demonstrated implantable neural probes with PICs that emit single beams, focused beams, light sheets, and steerable beams [30, 31, 40–43]; however, none had co-integrated electrophysiology recording capabilities, crucial for recording neural activity in deep brain regions. Fabrication on 200-mm diameter Si wafers in a commercial foundry ensures scalable manufacturing for broad dissemination in the future.

The application of these probes in *in vivo* experiments allowed us to compare the effects of different beam profiles on neural activity. Compared to probes that emitted low-divergence beams, the probes with light sheet emission that spatially distributed the optical power stimulated more neurons with minimal temperature increase. They also recorded more stimulated neurons and caused higher firing rate fatigue at lower output intensities. In the hippocampus, the light sheets elicited network-wide responses, typically necessitating high optical power outputs, triggering seizures in an epilepsy mouse model while keeping the temperature increase to less than 1°C to reduce the risk of tissue damage and thermally induced electrophysiological artifacts [11–13]. This work shows the potential of wafer-scale nanophotonic tools in probing deep brain network dynamics via electrophysiology and optogenetics, with customized optical beam profiles for enhanced spatial control over photostimulation.

## II. RESULTS

### A. Nanophotonic neural probe technology

The nanophotonic neural probe system is illustrated in Fig. 1. The probes were passive and used an off-chip laser source and recording electronics, minimizing the risk of tissue heating. Each probe was connected to an external laser scanning system and an electrophysiology data acquisition circuit board for simultaneous photostimulation and electrophysiology recording. The nanophotonic neural probes were fabricated on 200-mm diameter Si wafers using deep ultraviolet (DUV) lithography at Advanced Micro Foundry. Figures 2a and b show a photograph of the wafer and an illustration of its cross-section, respectively. The probe comprised a single layer of silicon nitride (SiN) for optical waveguides and 3 aluminum (Al) metal routing layers. Titanium nitride (TiN) was used to form biocompatible surface microelectrodes. The probe thickness could be reduced to 40 - 60 *µ*m through the foundry wafer backgrinding process followed by post-processing polishing, as shown in Fig. S1. Further details of probe fabrication are provided in Methods.

**FIG. 1.**
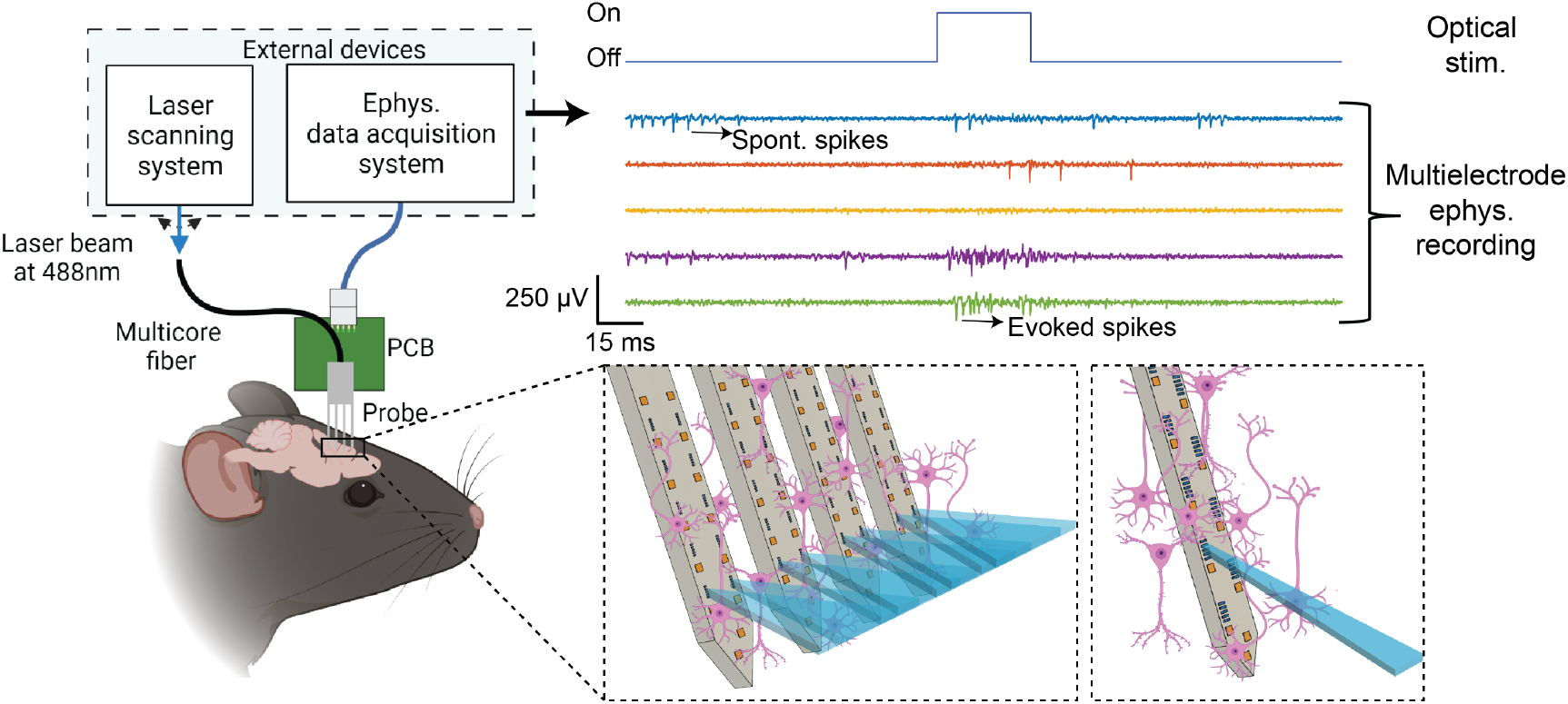
Conceptual illustration of the nanophotonic neural probe system. Various optical beam emission profiles, engineered by the grating design, can be directly generated in deep brain regions without any bulk optical components. A low-divergence beam is suitable for localized stimulation while a light sheet beam enables stimulation in a layer. The external laser scanning system addresses optical emission sites on the probe for multichannel photostimulation. The electrophysiology data acquisition system is used for recordings.

**FIG. 2.**
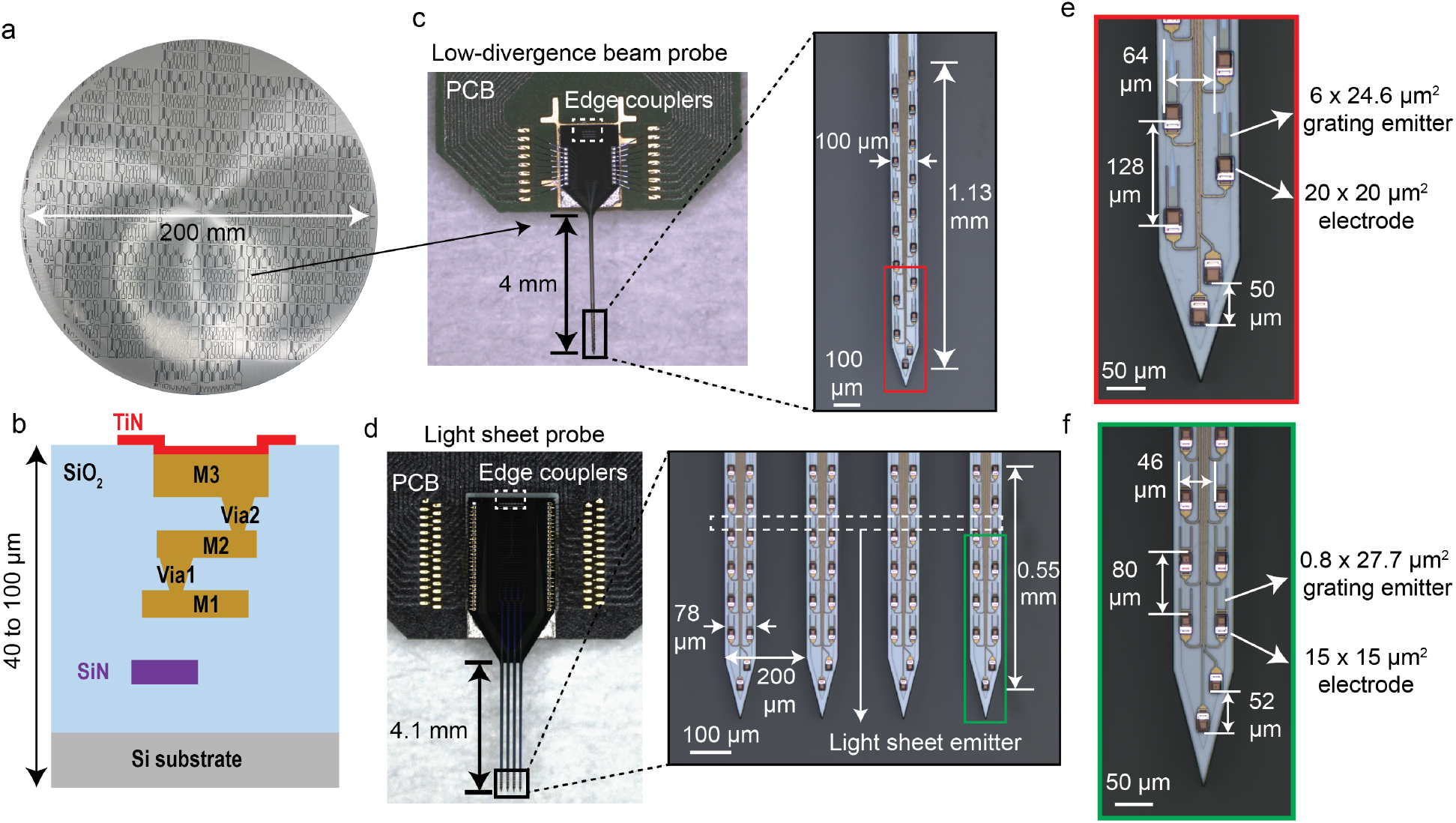
Overview of the neural probes. **a** 200-mm foundry-fabricated wafer with *>* 800 probes with various probe shapes and PIC designs and microelectrode arrangement. **b** Cross-sectional schematic of the hybrid neural probe platform. It supports one layer of SiN waveguide and 3 layers for metal routing, and TiN surface electrodes for electrophysiology recording. Photographs of **c** a low-divergence (LD) probe and **d** light-sheet (LS) probe wirebonded to printed circuit boards (PCBs) with insets showing optical micrographs of their shanks. Both probes have a shank length of *∼* 4mm. LD probes have a shank width of 100 *µ*m. The 18 electrodes and 16 emitters are distributed 1.13 mm at the tip of the shank. LS probes have 4 shanks each with a width of 78 *µ*m and a shank pitch of 200 *µ*m. 5 light sheet emitters and 56 electrodes (14 electrodes on each shank) are distributed 0.55 mm along the shanks. Optical micrographs of the shank on **e** an LD probe and **f** an LS probe at higher magnification. The LD probes feature electrodes and grating emitters with dimensions of 20 *×* 20 *µ*m^2^ and 6 *×* 24.6 *µ*m^2^, respectively. The vertical pitch between electrode and emitter pairs is 128 *µ*m, with a horizontal distance of 64 *µ*m between adjacent electrodes. The LS probe has electrodes and grating emitters with a size of 15 *×* 15 *µ*m^2^ and 0.8 *×* 27.7 *µ*m^2^, respectively. The vertical pitch between electrode and emitter pairs is 80 *µ*m, while the horizontal distance between two electrodes is 46 *µ*m. The scale bars are 100 *µ*m in **c** and **d** and 50 *µ*m in **e** and **f**.

To demonstrate the capability to tailor the beam emission profiles, we designed two types of probes with different gratings as shown in Fig. 2c-f. The first probe type, hereafter referred to as the “LD probe”, emitted low-divergence optical beams from a single shank. Each LD probe had 16 uniform gratings and 18 electrodes, each with a dimension of 20 *×* 20 *µ*m^2^, on a 4-mm long and 100 *µ*m wide shank. The shank length was capable of reaching a depth beyond the hippocampus in a mouse brain. The vertical and horizontal center-to-center distances of two adjacent electrodes were both 64 *µ*m, resulting in a detection distance of electrophysiology of 1.1 mm along the shank. The grating emitter had a grating pitch of 440 nm and a duty cycle of 50 % with dimensions of 6 *×* 24.6 *µ*m^2^, forming a low-divergence beam similar to the work presented in [40]. The vertical pitch of the electrodes and the emitters were both 128 *µ*m. The second probe type, hereafter referred to as the “LS probe”, emitted light sheets for layer-wise photostimulation. Each LS probe had four 4-mm long shanks and 5 light sheet emitters. The light sheets were formed by overlapping grating emissions from 8 grating emitters on the 4 shanks, and the design is described in [30]. Each light sheet grating emitter had dimensions of 0.8 *×* 27.7 *µ*m^2^, featuring a narrower grating width compared to the emitters on the LD probe. This design choice was aimed at achieving a larger beam divergence angle along the width of the shank to facilitate sheet synthesis. On each of the 4 shanks of the LS probe, there were 14 electrodes with dimensions of 15 *×* 15 *µ*m^2^ and spaced at a vertical pitch of 80 *µ*m. The light sheet emitters had the same vertical pitch as the electrodes of 80 *µ*m. The width of the shanks was 78 *µ*m, and the shank pitch was 200 *µ*m.

To support the photostimulation functionality of the probe, light from an external laser source was guided to the edge couplers on the chip with a custom 16-core single-mode multicore fiber (MCF) [44]. A microelectromechanical system (MEMS) mirror-based free-space laser scanning unit deflected a laser beam into the cores of the MCF, each of which was aligned to an edge coupler on the neural probe at the other end of the MCF. The laser scanning unit was similar to the one reported in [30, 31] with additional features to improve the stability of the free-space-to-fiber optical coupling efficiency for hours-long animal experiments (see Supplementary Section S2). The probe was wire-bonded to a printed circuit board (PCB) with Al or gold (Au) wire to connect to a commercially available Intan amplifier headstage and the Open Ephys data acquisition board for electrophysiology readout.

### B. Physical characterization of the neural probes

The as-fabricated TiN electrodes, with widths of 15 and 20 *µ*m, had respective impedances of 8.4 and 4.7 MΩ at 1 kHz, which were relatively high compared to the input impedance of the Intan amplifier (13 MΩ at 1 kHz). We developed a laser post-processing technique, described in Methods, to roughen the electrodes and reduce the impedance to *<* 2MΩ for recording [45]. Figures 3a and b show that the electrodes darken after post-processing, and scanning electron micrographs (SEMs) confirm that the electrode surface was intact but had a more textured surface following the laser treatment. Figure 3c shows the reduction in impedance. The mean impedance of the electrodes with widths of 15 (n = 18 electrodes on an LD probe) and 20 *µ*m (n = 54 electrodes on an LS probe) was reduced to 1.48 MΩ and 1.32 MΩ, respectively, with laser treatment. The electrode impedance decreased further to *<* 1 MΩ after immersing the probes in 1% Tergazyme solution for probe cleaning after animal experiments. The exact mechanisms for the impedance reduction have yet to be determined but may be due to the modification of the electrode surface in basic solutions (see Supplementary Section S3). The noise level of the raw electrical recording in 1*×* phosphate buffered saline solution (PBS) solution was 5.2 *±* 0.38 *µ*V and 4.2 *±* 0.23 *µ*V (root mean squared) for post-processed electrodes with widths of 15 and 20 *µ*m, respectively.

**FIG. 3.**
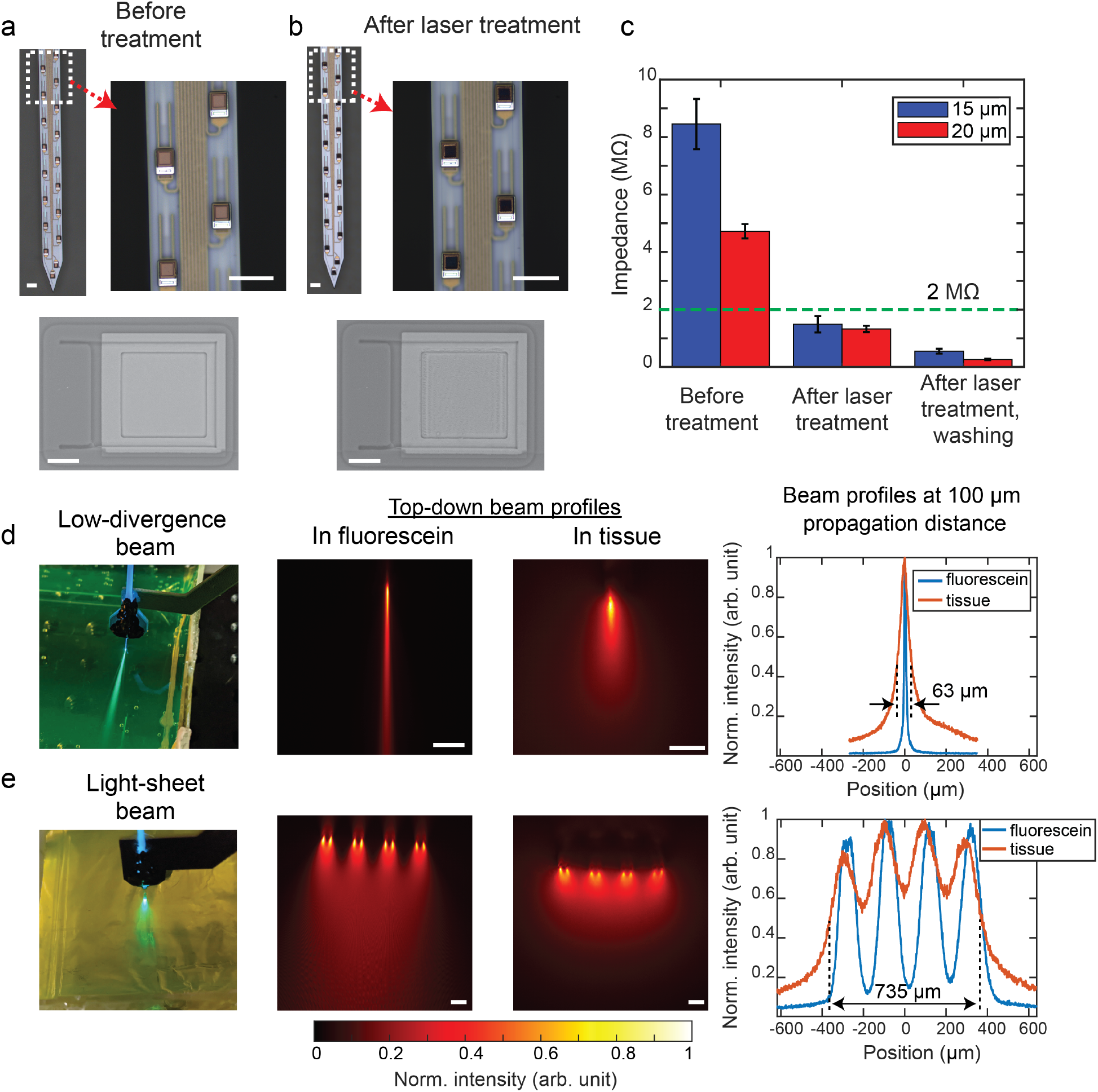
Characterization of the probe. Optical micrographs of the LD probe shank and scanning electron micrographs (SEMs) of an electrode on the LD probe before **a** and after **b** the laser treatment. The bottom SEMs show that the laser treatment created microstructures absent from the pretreatment condition. The scale bars are 50 *µ*m in the micrographs of the shank and 20 *µ*m in the SEMs of the electrode. **c** Comparison of impedances before and after laser treatment of an LD probe with 20 *µ*m electrodes and an LS probe with 15 *µ*m electrodes. The green dashed line shows the 2 MΩ limit to achieve a reasonable signal-to-noise ratio (SNR) for electrophysiology recording. The impedance reduced to *<* 1 MΩ after the Tergazyme wash. The error bars represent the standard deviations (SD). Top-down view of **d** the low-divergence beam and **e** the light sheet beam propagating in fluorescein solution and in fluorescein-stained cortical brain slices. Line profiles for both types of beams in fluorescein and tissue at a propagation distance of 100 *µ*m are plotted to illustrate the difference in beam size. The scale bars in the beam profile images are 100 *µ*m.

Regarding the optical properties of the probes, the insertion loss before and after the MCF-to-chip attachment process is discussed in the Supplementary Section S4. Each emitter in the packaged probes could provide *>* 5 *µ*W of optical power, equivalent to an output intensity of 64.2 mW/mm^2^ for the LD probe and 3.9 mW/mm^2^ for each of the 8 gratings on the LS probe, exceeding the optical threshold intensity to stimulate ChR2 neurons [1]. These output intensities were estimated using the beam size (1*/e*^2^ width) calculated from finite-difference time-domain (FDTD) simulations presented in Fig. S5b. The lower output intensity of the LS probe is the result of the optical power being distributed over 8 grating emitters and the wider beam divergence angle.

Figure 3 d and e show the emission patterns of the two probe types in non-scattering (fluorescein solution) and scattering (fluorescein-stained cortical mouse brain slices) media. For the LD probe, the full-width at half-maximum (FWHM) beam width at a propagation distance of 100 *µ*m in fluorescein was 10 *µ*m. Over this distance, for the LS probe, the light sheet beam profile was not yet fully formed. Tissue scattering caused the FWHM of the low-divergence beam to widen to 63 *µ*m, while the 4 lobes of the LS probe partially merged into a sheet with an FWHM of 735 *µ*m. In general, the light sheet was 11 times wider than the low-divergence beam in tissue. The output power distribution across multiple shanks on the LS probe can lead to a wider stimulation region at lower output intensities, as shown in Fig. S6a. Supplementary Section S5 describes the details of the estimated beam intensity profile of the two beam types.

The side-view beam profile measured in fluorescein solution shows that each emitter can be independently addressed with no visible crosstalk between emitters (Fig. S7 and Supplementary video 1). Our previous work in [44] also demonstrated that the channel crosstalk at *λ <* 532 nm in a 1-m long MCF was, in the worst case, *< −*35 dB, lower than the crosstalk in other optical switching mechanisms demonstrated in nanophotonic neural probes (i.e., wavelength division multiplexing or MZI optical switches of *<* -13 dB) [26, 28]. This ultralow crosstalk is critical in enabling our probes to output confined beam profiles from each emitter under high-power settings (*>* 100 *µ*W).

### C. Spatially selective optogenetic stimulation

To assess the optogenetic stimulation and electrophysiology functionalities of the probes, we conducted *in vivo* experiments in awake head-fixed Thy1-ChR2-eYFP mice (see Methods for experimental procedures). A total of 6 mice were used for the LD probe and 7 mice for the LS probe. The probes were implanted in layers V and VI between the somatosensory and motor cortex (AP: -0.5 mm, ML: 1.2 mm) as shown in Fig. 4a (histological confirmations are presented in Fig. S8a). Our photostimulation pattern was at a wavelength of 488 nm and consisted of a 2-s long train of 30-ms optical pulses at a repetition rate of 5 Hz. We repeated the pulse train 6 times with 20 s resting time in between the pulse train, resulting in a total of 60 pulses. To estimate the effective stimulation area of each beam profile, we conducted trials with different power levels and grating emitters on each probe. As shown in Fig. 4b, example traces of the evoked spiking response with a 30-ms pulse from the LS probe demonstrate that Sheet 1 and 5 can selectively activate neurons at two different depths. The same stimulation pattern at higher optical power did not cause spiking responses in the control experiment conducted with an anesthetized wild-type mouse, as presented in Fig. S9a. Figure 4c shows the waveforms of two activated neurons and their corresponding autocorrelograms during the stimulation trial, which showed a low violation of the refractory period. Furthermore, the mean firing rate of the two neurons during the 60 repetitions of the 30-ms pulses, i.e., induced by the light sheet (in Fig. 4d), returned to a near-baseline level if the photostimulation was 240 *µ*m away (equivalent to 3 emitters) from the location with the highest evoked firing rate. This observation underscores the localized confinement of the thickness of the sheet. An example demonstrating spatially selective photostimulation for the LD probe is shown in Fig. S10.

**FIG. 4.**
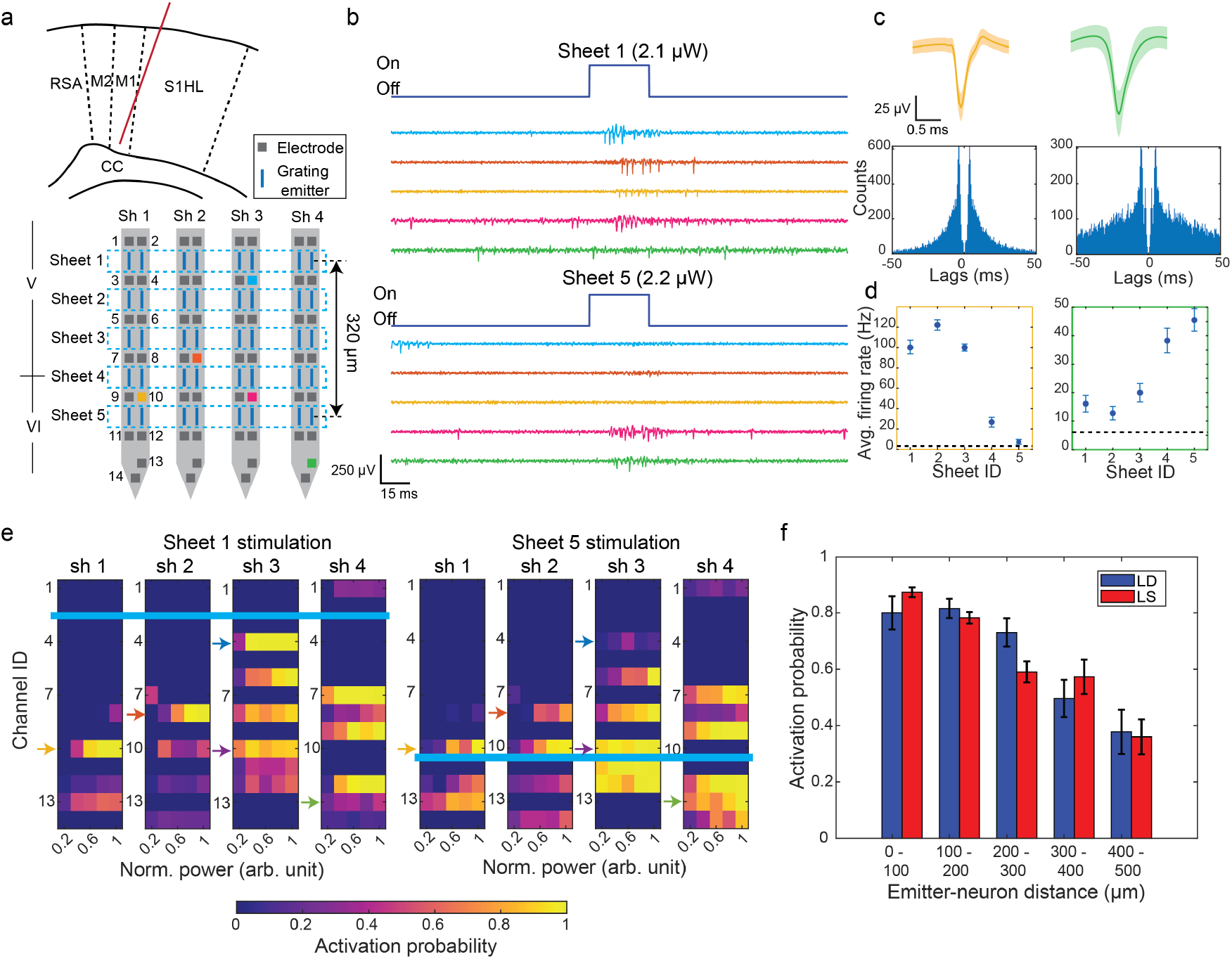
Demonstration of spatially selective optogenetic stimulation with an LS probe in *in vivo* awake head-fixed mice. **a** The implanted trajectory of the LS neural probe was between the motor cortex and the somatosensory cortex. All electrodes and light sheet emitters were in cortical layers V and VI. Channel IDs of the electrodes on the shank and sheet emitter numbers are labeled. The distance between Sheet 1 and 5 is 320 *µ*m. **b** Snapshots of the filtered voltage traces (common average referencing (CAR) + bandpass filtered (300-6000 Hz) + artifact subtraction) from 5 electrodes showing evoked spikes during the 30 ms long optical pulse from Sheet 1 and Sheet 5 emitters. The optical power outputs of the sheets were 2.1 and 2.2 *µ*W, respectively. The colors of the traces match the colors of the electrodes in **a**. **c** Waveforms of two example single units and their autocorrelograms. The color of the waveform matches the color of the recorded traces. The solid lines represent the mean waveform and the shaded area represents the standard deviation (SD). **d** Mean firing rate of the two units during the 30 ms optical stimulation from the 5 light sheet emitters (*n* = 60 pulses). The error bars denote the standard error of the mean (s.e.m). The dashed lines represent the mean baseline firing rate, averaged over 10 seconds immediately preceding each stimulation pulse train. The output power of the 5 sheets ranged from 2.1 to 2.5 *µ*W. **e** Mean ActProb of the neurons on the 4 shanks with increasing optical powers of Sheet 1 and Sheet 5. The power is normalized to the trial with the maximum power output (Sheet 1: 3.5 *µ*W, Sheet 5: 5.7 *µ*W). The color-coded arrows indicate the position of the example traces in **b** and the blue lines show the positions of the light-sheet emitters. **f** The mean ActProb versus emitter-to-neuron distance along the shank of the LD and LS neural probes for all animal trials. The plot includes neurons that exhibited an ActProb *>* 80% after stimulation by any emitter with an output power ranging between 2.5 to 6 *µ*W. The error bars denote s.e.m.

As a measure of the extent of the brain region that was stimulated, we define a metric, activation probability (ActProb), as the probability of detecting a spike within the 30-ms optical pulse for each neuron. Figure 4e shows an example spatial heatmap of ActProb across the 4 shanks for photostimulation using Sheet 1 and 5 on the LS probe. For electrodes detecting more than one neuron, we report the multi-unit ActProb. Here, we observed that both Sheet 1 and 5 could evoke spiking activity with ActProb *>* 90% across the four shanks, indicating sufficient emission intensity from the 8 grating emitters that constitute the light sheet emitter.

Figure 4f shows the mean ActProb for all animal trials as a function of the optical emitter-to-neuron distance measured along the shank. At least 4 emitters were used for each animal trial to observe the stimulation response in the cortex. To only consider neurons with stimulated responses, the plot included neurons with an ActProb *>* 80 % following stimulation by any emitter with an output power ranging from 2.5 to 6 *µ*W. Both the light sheet and the low-divergence beam repeatedly elicited spiking activity (mean ActProb *>* 75%) within 200 *µ*m of an emitter while the mean ActProb for both types of probe decreased to *<* 40% more than 400 *µ*m from the emitter. There is no statistically significant difference in ActProb (p *>* 0.05) between the two probe types across all emitter-to-neuron distance intervals, suggesting that both beam types have a similar stimulation extent along the probe shank. This outcome can be attributed to the similar beam thickness of both beam profiles, as demonstrated in the simulated beam profile presented in Fig. S5. However, for optical output powers in the range of 2.5-6 *µ*W (equivalent to output intensities of 32.1-77.1 mW/mm^2^ for the low-divergence beam and 2-4.7 mW/mm^2^ for the light sheet beam, as estimated with the simulated beam size), the LS probe generally activated more neurons with ActProb *>* 80 %, an indication of reliable stimulation [46], than the LD probe. This is likely due to the wider optical emission profile and the larger number of electrodes on the LS probe. Of the recordings captured by the LS probe, 69 of 154 single units from 7 mice had a high ActProb *>* 80%. In contrast, with the LD probe, only 22 of 49 single units recorded from 6 mice had ActProb *>* 80%.

**FIG. 5.**
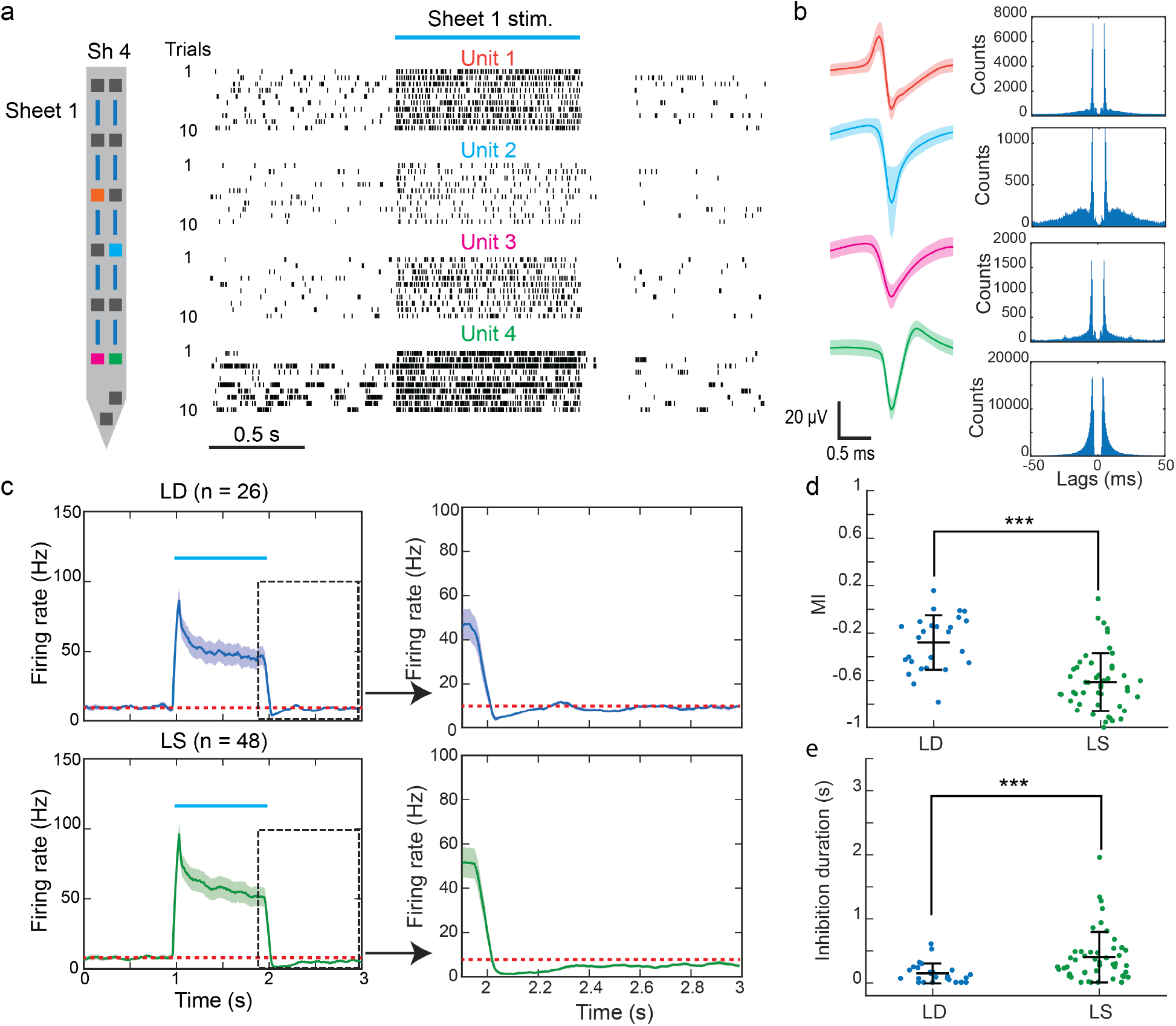
Evoked spiking response in layer V and VI of motor and somatosensory cortex with 1-s CW stimulation. **a** Spiking raster plot of 4 example units during the 1-s CW stimulation period over 10 trials. All 4 units were recorded from shank 4 of an LS probe. The blue solid line above the raster plots indicates the stimulation time. The position of the single units is color-coded on the shank. **b** The waveforms and autocorrelograms of the 4 example units. **c** Mean PSTHs of all activated units (increase in firing rate *>* 5 Hz during stimulation and *>* 3 Hz pre-stimulus firing rate) for both probe types recorded in all animal trials. 26 single units were recorded by the LD probe and 48 single units by the LS probe. The shaded region represents the s.e.m. of the trace. The red dashed lines indicate the mean pre-stimulus firing rate calculated over the 10 seconds preceding the stimulation, and the blue line labels the stimulation period. The insets show a zoom-in PSTH for post-stimulation response. Both probes can induce firing rate fatigue while, on average, evoked units by the LS probes took longer to recover to 80% of the pre-stimulus firing rate (LS: 880 ms, LD: 130 ms). **d** Modulation index and **e** inhibition duration of each stimulated unit evoked by LD and LS probes. The center horizontal lines and the error bars in d-e indicate mean and SD, respectively. *** denotes *p <* 0.001.

**FIG. 6.**
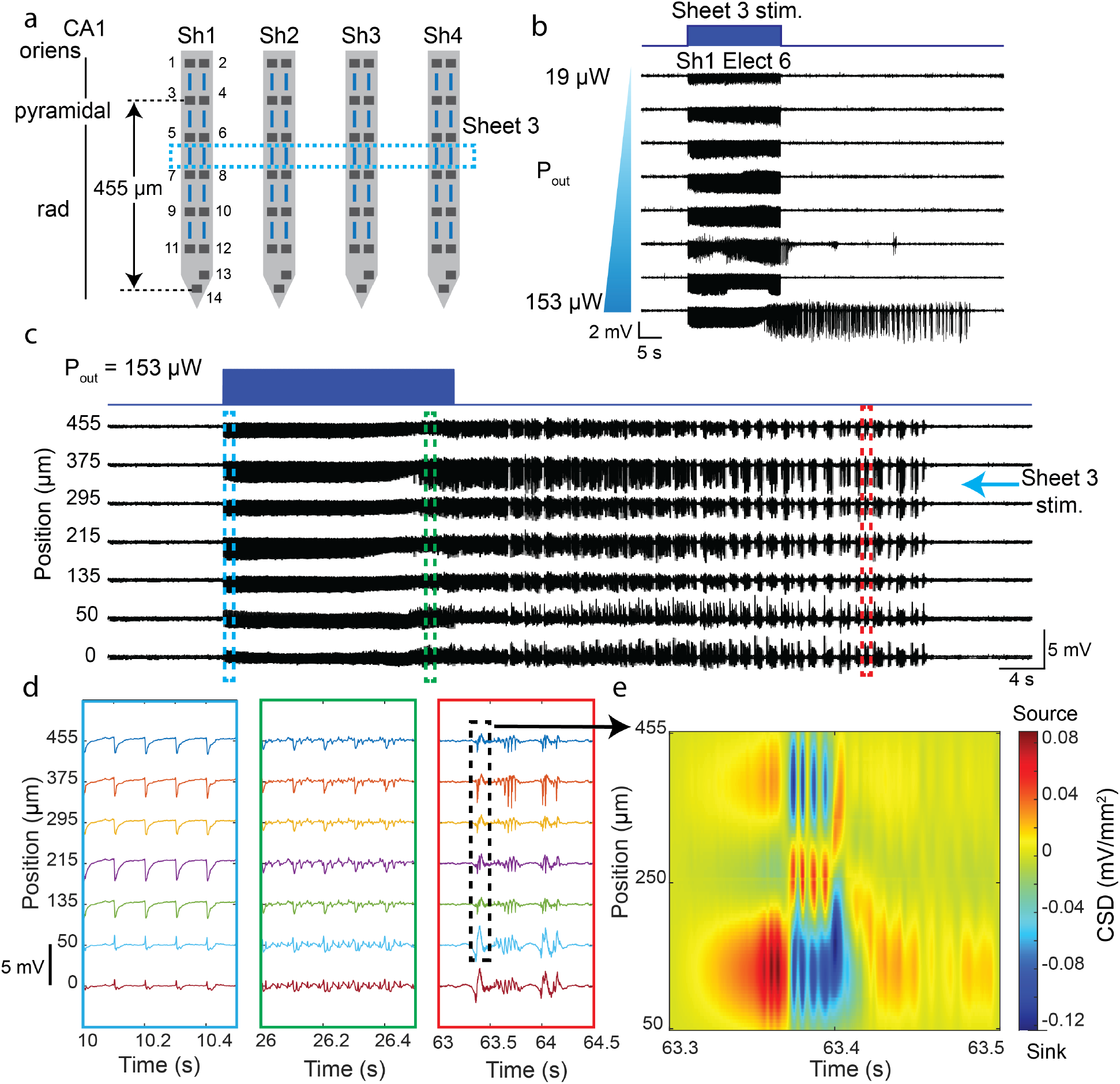
Optogenetically induced seizure by an LS probe in CA1 of the hippocampus in an epilepsy mouse model. **a** Insertion position of the probe in the hippocampus. Channel IDs of the electrodes on the shank and the sheet emitter numbers are labeled. Sheet 3 was used to optogenetically elicit seizure. The vertical distance from Electrode 3 to Electrode 14 was 455 *µ*m, the region where the preceding analysis was performed. **b** Evoked LFP response (CAR + bandpass filtered (5 - 300 Hz)) recorded on Electrode 6 on Shank 1 across different optical power trials. **c** Evoked LFP responses from CA1 along the right side of Shank 1 during a successful seizure induction trial. **d** Snapshots of the LFP trace in three time intervals during the time traces of the seizure development in **c**. **e** Current source density (CSD) map along Shank 1 at the onset of one of the after-discharges. Electrode 14 is excluded from the CSD calculation due to its horizontal offset with respect to other electrodes on the right side of Shank 1.

### D. Firing rate fatigue induced by continuous optogenetic stimulation

In the same animal trials described in the previous section (LD: 6 mice, LS: 7 mice), we also investigated the evoked spiking response from 1-s of continuous-wave (CW) stimulation. In each trial, we applied 1-s long stimulation 10 times with 30 s of rest between each stimulation. The stimulation pattern was repeated for at least two light emitters on the probe. Figure 5a shows the raster plots of four example units under 1-s CW stimulation by an LS probe in one of the animal experiments (their spike waveforms and correlograms are presented Fig. 5b). During stimulation, the evoked units exhibited a high mean firing rate of *>* 20 Hz. Furthermore, a brief inhibition period (*<* 1 s) occurred in all four units after the 1-s stimulation. The firing rate of 500 ms after the stimulation for all 4 units was statistically lower (p *<* 0.01, one-tailed Wilcoxon signed-rank test) than the firing rate 10 s prior to the stimulation. This inhibition is possibly due to firing rate fatigue [47], as the inhibition response was weaker with low-frequency stimulation (see Fig. S11).

To investigate the average response of the stimulated neurons to the 1-s photostimulation, Fig. 5c shows the mean peristimulus time histogram (PSTH) of all photostimulated units in the animal trials for both probe types. The stimulated units were taken to be those meeting the following three criteria: 1) a pre-stimulus firing rate *>* 3 Hz measured at 10 s before stimulation, 2) the pre-stimulus firing rate is statistically lower than the firing rate during the 1-s stimulation (n= 10 trials, p *<* 0.05, one-tailed Wilcoxon signed-rank test), and 3) an average increase in firing rate of *>* 5 Hz during the 1-s stimulation compared to the pre-stimulus firing rate. Here, we observed a strong but decaying firing rate during the 1-s photostimulation period for both probe types. The inhibition response also occurred after the 1-s stimulation. However, the inhibition response appeared stronger for the LS stimulation, as evidenced by the longer time required to recover to the pre-stimulus firing rate. Defining the inhibition duration as the time it takes the neuron to recover to 80% of the mean pre-stimulus firing rate, the inhibition duration of the LD probe trials was 130 ms and the LS probe trials was 880 ms. To evaluate whether the difference was statistically significant, we analyzed the strength and duration of the inhibition response for the stimulated units. The strength of the inhibition response was quantified using the modulation index, MI, defined As

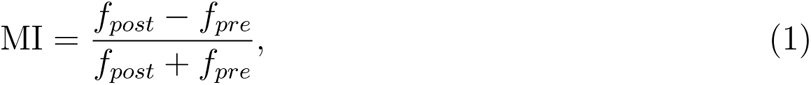

where *f_pre_* is the firing rate 10 s before the stimulation, *f_post_* is the firing rate 0.5 s after stimulation. The MI metric was used in [48] to quantify photostimulated responses. As presented in Figs. 5d and e, both metrics suggest that the LS probe statistically elicited stronger inhibition responses than the LD probe. Further categorizing the trials into low- and high-power settings for the LS and LD probes (Supplementary Section S11) shows that the LD probe at higher output power could reduce the level of statistical significance in the difference in inhibition strength with *∼* 11 *×* higher peak output intensities (see Supplementary Section S11 for intensity calculations). This experiment provides additional evidence that the LS probe could elicit stronger responses than the LD probe with lower output intensities.

### E. On-demand optogenetically induced seizure

The larger spatial extent of the LS beam profile resulted in a reduced output intensity at a particular output power, leading to a lower temperature increase. As an example of how this can be applied to network-wide effects, we used an LS probe to induce seizures in an optogenetic epilepsy mouse model. The model was Thy1-ChR2-eYFP mice injected with Kainic acid to increase brain excitability [49] (see Methods for details on the preparation). An LS probe was inserted into CA1 of the hippocampus, as illustrated in Fig. 6. For the stimulation pattern, we applied a pulse train with a 5-ms pulse width at 10 Hz, lasting 20 seconds, as shorter duty cycles have been shown to increase the probability of optogenetically-induced seizures [50]. To determine the power threshold to induce seizures, we repeated the stimulation patterns with varying optical power levels starting from 19 *µ*W and increasing at a step size of 16-21 *µ*W until a seizure was induced.

As shown in Fig. 6b, a range of optical output powers caused evoked potentials at the onset of the optical pulse train; however, only an optical power output of 153 *µ*W led to a sustained after-discharge lasting approximately 40 s, indicative of a seizure. The expected temperature increase, which was only due to optical absorption in tissue, was estimated to be *<* 1 °C. Supplementary Section S12 shows that the temperature would increase from 37°C to a maximum of 37.94°C for a CW output power of 280 *µ*W from a light sheet emitter.

Figures 6c and d show the spatial distribution of the local field potential (LFP) in one of the shanks when a seizure was induced. The seizure could be classified into three distinct stages similar to the pattern observed in [50]. First, a combination of light-induced artifacts and time-locked evoked potentials were observed only at the onset of the optical pulse. Second, the LFP exhibited a combination of time-locked responses and spurious after-discharges. Third, after stimulation, self-sustained after-discharge responses with high amplitude persisted for approximately 40 seconds. The behavior exhibited by the mouse during the third stage corresponded to a seizure severity with a Racine scale of 5 [51]. We used the probe to measure the spatial distribution of the LFPs, allowing us to identify the sink and source relationships between the channels, as shown in Fig. 6e. Overall, this experiment validates the ability of the nanophotonic neural probe to deliver sufficient optical power to optogenetically induce seizures on demand with minimal heating and opens the possibility of investigating the dynamics of seizures across regions of the brain with a single probe device [50].

## III. DISCUSSION

In this work, we have reported nanophotonic neural probes with capabilities for electrophysiology recording and photostimulation using flexibly designed light emission patterns. Our neural probes demonstrated advantages that had been lacking in other implantable probes with combined electrophysiology and optogenetic stimulation functionalities. Compared to other nanophotonic neural probes with SiN waveguides and microelectrodes, the probes here have the most integrated optical emitters (16 emitters) that deliver optical beam profiles beyond those of single gratings while offering the lowest optical crosstalk (*<* -35 dB) using the spatial multiplexing scheme [26, 28, 29]. Low optical crosstalk is particularly critical in high-power output applications. For example, with 100 *µ*W of output power, equivalent to 78.6 mW/mm^2^ from the LS probe, *< −*20 dB of crosstalk is desirable to ensure that only the main output emission elicits spikes. While optoelectronic probes with integrated µ-LEDs and µ-OLEDs [24, 25] have high densities of light emitters (*>* 100 channels), they are not well suited to deliver high optical output powers to address a large population of neurons due to their low electrical-to-optical power conversion efficiencies (*<* 3%) and heat dissipation. For example, although the µ-LED probe can provide relatively high output power at high current injection (53 *µ*W output at 13 mA), a 10 ms-long optical pulse would already cause a device temperature rise of 1 °C as discussed in [21]. The thermal limit may prohibit the use of optoelectronic probes to stimulate network-wide activity or neurons that are further away from the probe. Fiber-based approaches (i.e., tapered fiber and thinned fiber matrix packaged with Si microelectrode probe [16, 52]) can deliver high optical powers to stimulate large populations of neurons, but they lack the scalability to realize high-density electrode and optical emitter arrays on a single implantable device. Table IV compares different Si neural probe technologies with photostimulation capabilities.

We envision several directions for our nanophotonics-based neural probes. First, with our current approach, it is difficult for the neural probes to scale to a high optical emitter channel count, say beyond 100, due to the MCF spatial multiplexing scheme (i.e., the manufacturing of the MCF and precise alignment of the MCF to the waveguides on chip). To increase the number of optical channels, one approach is to incorporate PICs in the base of the probe for switching and wavelength division multiplexing. For example, 1152 optical emission sites can be realized using 16 fiber cores as inputs, each coupled to a 1*×* 8 optical switch, and each switch output is connected to a 1 *×* 9 wavelength division multiplexer. Such a channel count is similar to the state-of-the-art µ-OLED neural probe while offering an output intensity at least one order of magnitude higher for each emitter (*>* 5 mW/mm^2^ vs. 0.25 mW/mm^2^). Another limitation of the current device is that, with electrodes and emitters integrated on the same probe, only evoked responses in close proximity to the probe can be detected; thus, a probe cannot fully capture the effects of LS and LD beam stimulation. A solution is to implant several probes to extend the detection horizon. Another promising and more compact method, as demonstrated in [53], is to stack multiple probes using 3D integration approaches to enable high-density brain interrogation within a smaller volume. Lastly, to simplify the fiber-to-chip packaging, we are working toward laser integration while managing the thermal load on the probe.

In summary, we have demonstrated implantable neural probes with capabilities for simultaneous electrophysiological recording and patterned photostimulation. Unique to these probes is the use of integrated nanophotonics to achieve custom and tailored light emission patterns. Probes emitting low-divergence beams and light sheets enabled localized and network-wide optogenetic stimulation, respectively. Since the light source was not integrated on-chip, the probes could deliver high optical intensities with minimal heat dissipation. Light sheet emission was found to elicit a higher level of activity at lower output intensities compared to the low-divergence beam, as evidenced by the number of neurons stimulated with high ActProb and stronger firing fatigue, and could induce seizures in an epilepsy model while keeping the expected temperature increase at *<* 1 °C. The wafer-scale foundry fabrication of these probes is a significant step toward broad dissemination of the technology to the neuroscience community. Ongoing work on expanding the functionalities of these neural probes, such as increasing the number of optical channels, integrating power-monitoring photodetectors and temperature sensors, as well as incorporating microfluidic channels for chemical sensing and drug delivery, is currently underway. Foundry Si photonics technology promises to enable a new generation of multifunctional neural implants for multimodal neural stimulation and recording.

### IV. METHODS

### A. Neural probe fabrication

The neural probes were manufactured on 200-mm diameter Si wafers at Advanced Micro Foundry (AMF). First, a layer of silicon dioxide (SiO_2_) was deposited on the Si wafer followed by the deposition of SiN for the waveguide core. The photonic layer can be either 120-nm thick low-pressure chemical vapour deposition (LPCVD) or 200-nm thick plasma-enhanced chemical vapour deposition (PECVD) SiN. 193-nm DUV photolithography and reactive ion etching (RIE) were used to define the waveguide pattern. Then a top layer of SiO_2_ was deposited on the waveguide. To enable electrophysiology capability, 3 layers of Al routing metallization with vias connecting between the layers and TiN surface microelectrodes were subsequently formed. Chemical mechanical polishing (CMP) was used for layer planarization. The outline of the probes and the edge coupler facets were formed with deep trench etching. Finally, the probes were separated from the wafer by thinning the wafer thickness to approximately 100 *µ*m through backgrinding. Additional mechanical polishing was performed on some singulated probes using a fiber polisher (NOVA Optical Polishing System, Krell Technologies, Neptune City, NJ, USA) as a post-processing step to thin the probes to 40-60 *µ*m as illustrated in Fig. S1

### B. Laser treatment of TiN electrodes

A laser treatment process was developed to roughen the electrodes to reduce their impedance to below 2 MΩ. To control the laser scanning pattern on the micrometer-sized electrode, we performed the laser roughening process using a 2-photon microscope (Bruker Instruments). A femtosecond laser that emits light with a 10 kHz repetition rate and a wavelength of 1035 nm (Coherent Monaco) was utilized for the roughening process. For sample preparation, we placed the probe on a glass slide, covering it with a glass coverslip to hold it in place. Distilled water was added on top of the coverslip so the sample can be imaged with the water immersion objective on the microscope (16*×* Nikon CFI LWD plan fluorite objective). Once we located the microelectrodes under the microscope, we applied a 2 dimensional (2D) array of spiral patterns to form a rectangular shape slightly smaller than the electrode’s dimensions to prevent chipping of the electrode edges during the laser process. The scanning pattern was controlled with a Galvo mirror. The mean incident power from the microscope objective is 60 - 80 *µ*W with a pulse duration of 270 fs and a spot size of *≈* 1.2 *µ*m in diameter (estimated with a diffraction-limited spot size as the electrode is in focus). A single scan across a 15 *×* 15 *µ*m^2^ electrode takes 0.128 s and a 20 *×* 20 *µ*m^2^ electrode for 0.2 s. We repeated the scanning pattern 8 to 32 times for each electrode. The electrodes were inspected with an optical microscope, confirming two criteria: 1) the electrode surface exhibited a darkened appearance, and 2) the electrode remained intact.

### C. Electrode impedance measurement

To measure the impedance of the electrodes, the probe shanks were immersed in 1*×* PBS with an Ag / AgCl wire as the ground electrode. Three impedance measurement systems were used to characterize the electrode at different stages of the probe. For bare Si probes, we connected to the contact pads with a micro-needle and measured the impedance with an impedance analyzer (Keysight E4990A). The voltage input was set at 10 mV to avoid electrolysis in water. For probes wirebonded to PCBs, an impedance meter (Nanoz, Whilte Matter LLC, Seattle, WA, USA) or an amplifier headstage (Intan Technologies, Los Angeles, CA, USA) with an electrophysiology data acquisition system (Open Ephys) was used to measure the impedance in PBS. All reported impedances were characterized at 1 kHz.

### D. Neural probe packaging

The neural probes were attached to PCBs and custom visible-light 16-core single-mode MCFs (Corning, Corning, NY, USA). Each probe was attached to the PCB with a metal substrate using heat-curable metal epoxy (Ablebond 84-1LMIT1, Loctite, Stamford, CT, USA), and bond pads on the probe were wirebonded to the PCB with either Au or Al wires. To insulate the wirebond, UV-curable encapsulation epoxy (Katiobond GE680, Delo, Germany) was applied and cured with a UV LED system (CS2010, Tholabs, Newton, NJ, USA). A layer of optically opaque epoxy (EPO-TEK-320, Epoxy Technology, Billerica, MA, USA) was subsequently applied to block wirebond from stray light, which can cause light-induced artifacts during recording.

Fiber-to-chip attachment was performed after the electrical packaging steps, as it required more precise alignment. The MCF was actively aligned to the edge couplers on the chip with a 5-axis piezoelectric actuated fiber alignment stage. To assist the fiber-to-chip alignment process, a top-down microscope and a photodetector were employed for power monitoring. Once optimal alignment was achieved, UV curable low-shrinkage epoxies (OP-67-LS and OP-4-20632, Dymax Co., Torrington, CT, USA) were applied incrementally to cover both the fiber and the chip, minimizing the alignment drift caused by epoxy shrinkage during curing. To provide stress relief, a 5-minute epoxy (Loctite, Westlake, OH, USA) was applied to the back of the fiber and PCB. Finally, an additional optically opaque epoxy encapsulated the fiber and the probe base, preventing stray light from reaching the brain. The packaged probe was placed on an optical table for a minimum of 12 hours to ensure complete curing of the epoxy before use in animal experiments.

### E. Simulations of the beam emission profiles

We simulated the beam emission profiles of the low-divergence beam and the light sheet beam in a scattering medium with the beam propagation method (BPM) similar to the method described in [31]. The scattering effect was accounted for in the simulation with consecutive planes of random phase mask with small variance along the propagation direction. We designed the phase masks, using the approach detailed in [54], and set the scattering coefficient (*µ_s_*) to 200 cm*^−^*^1^, the anisotropy of the tissue (*g*) to 0.83, and the attenuation coefficient (*µ_a_*) to 0.62 cm*^−^*^1^ [55].

The near-field optical emission profile was determined through finite-difference time-domain (FDTD) simulations. As the input light to the chip was depolarized to mitigate the power fluctuation of the scanning system due to polarization fluctuations in the 1 m long fiber, as detailed in Supplementary Section S2, we simulated the beam emission patterns for both transverse electric (TE) and transverse magnetic (TM) polarizations. The depolarized output light was represented by superimposing the TE and TM emission patterns generated by the grating emitter, and their intensity ratio was adjusted based on the experimental output coupling efficiency for each polarization. The electromagnetic field at 2 *µ*m above the oxide cladding (equivalent to *∼* 8 *µ*m above the grating emitter) was used as the input field for the BPM simulation. We computed the beam emission profile for a propagation distance of 200 *µ*m due to memory constraints of the workstation. For the light sheet emitter, only a single grating was simulated. The volumetric beam profile of a single beam was then replicated 8 times with a lateral offset that matched the design of the light sheet emitter array. The intensity of each emitter was corrected to account for the excess loss introduced by the extra crossings in the routing network. This power distribution was estimated using the mean of the highest 1% intensity measured at the output of each grating in fluorescein solution. We also rotated the volumetric beam profile to align the beam propagation axis with the horizontal axis to allow a top-down cross-sectional view to be extracted.

### F. Characterization of the beam emission profiles

To study the properties of different beam types, we inserted the probes in 1) fluorescein solution (non-scattering medium) and 2) fluorescein-stained cortical slices (scattering medium). A 10 *µ*M fluorescein solution was prepared with the fluorescein sodium salt powder (46960-100G-F, Sigma-Aldrich, Burlington, MA, USA) diluted in Milli-Q water with the pH controlled to be *>*9 using NaOH. The packaged probe was fixed to a metal rod mounted on a 4-axis micromanipulator (QUAD, Sutter Instrument Company, Novato, CA, USA). The probe was immersed in the solution at an angle such that the optical beam propagated parallel to the surface of the solution. An epifluorescence microscope was employed to capture the top-down view of the beam profile, while a second microscope was positioned horizontally to capture the side-view beam profile. To obtain an optically clear side view, a coverslip window was constructed on the side of the solution container. Both microscopes were equipped with GFP filters to reject the excitation wavelength.

For the tissue sample preparation, we prepared 1.5-2 % paraformaldehyde (PFA) fixed whole brains from wild-type mice (C57BL / 6J). The animal was initially anesthetized with isoflurane inhalation. The mouse was then transcardially perfused with 1*×* PBS followed by paraformaldehyde (PFA) (4%). The brain was then extracted and immersed in 1.5-2 % PFA at 4°C for 8 hours before being transferred to PBS solution for storage.

After fixing the brain, it was cut into slices with a thickness of 2 mm. The brain slices were then permeabilized in a 0.3% Triton X solution (Sigma Aldrich, Burlington, MA, USA) for 30 minutes, followed by three rounds of 5-minute washes in Milli-Q water. Subsequently, the slices were immersed in a 100 *µ*M fluorescein solution for 24-48 hours. Before use, the slices underwent another three rounds of 5-minute washes in Milli-Q water and were affixed to a petri dish filled with 1*×* PBS.

For beam profile imaging in tissue, an epifluorescence microscope was used to obtain the top-down beam profile from the gratings. The probe was positioned at an angle to ensure that the beam propagated parallel to the tissue surface. Typically, the probe was inserted into layers V and VI of the cortex, chosen for their relevance to *in vivo* experiments. After confirming that the emitter was inserted into the tissue, the probe was slowly retracted with a step size of 10-30 *µ*m while imaging the beam profile of the emitter. Once an emitter was outside the brain slice, the next emitter was addressed, and the imaging procedure was repeated. This iterative process helped minimize the impact of tissue scattering in depth on the beam profile.

### G. Animals

All experimental procedures described here were reviewed and approved by the animal care committees of the University Health Network in accordance with the Canadian Council on Animal Care guidelines. Adult male and female Thy1-ChR2-YFP mice (The Jackson Laboratory, Bar Harbor, Maine, stock number 007612) and C57BL/6J mice (Charles River Laboratories, Wilmington, Massachusetts) were kept in a vivarium maintained at 22 °C with a 12-h light on/off cycle. Food and water were available *ad libitum*.

### H. Experimental protocol for awake head-fixed animal experiments

Two to five days prior to the experiment, the head plate was mounted on the mouse skull with the following procedure. A Thy1-ChR2-YFP transgenic mouse or a wild type (C57BL/6J) mouse, at age over 60 postnatal days (P60), was anesthetized by induction with 5% and maintenance with 1% to 2% isoflurane/oxygen anesthetic and secured in a stereotaxic frame via ear bars (Model 902, David Kopf Instrument, Tujunga, California). We adopted a ground and reference configuration similar to that described in [56]. We drilled two small holes in the skull with a dental drill for the implantation of bone screws (Item No. 19010-10, Fine Science Tools), which served as ground and reference electrodes. One hole was located toward the front of the bregma on the ipsilateral side of the probe, and the other hole was toward the cerebellum on the contralateral side of the probe. The desired probe insertion position was also marked with a dental drill (stereotaxic coordinates of AP: 0 to -0.5 mm, ML: 1.2 mm for targeting motor and somatosensory cortex), as the head plate can obstruct the view of the bregma. The head plate was attached to the skull with dental cement (C&B Metabond, Parkell, Edgewood, NY, USA). After surgery, the mouse was placed back in the cage for recovery.

On the day of the experiment, the mouse was anesthetized following the same procedure and placed inside the stereotaxic frame. A circular craniotomy with a diameter of 1 - 2 mm was performed centered on the previously marked position for probe insertion. Typically, the dura was left intact to minimize possible motion artifacts caused by brain pulsation. For older mice (*>* P120), we sometimes remove the dura for ease of probe insertion. After the skull was removed, the mouse was transferred to an epifluorescence microscope. Its body was positioned within a 3D-printed cone to restrict body movements, while the head plate was attached to metal bars. Bone screws were connected to the ground and reference wires, which were soldered on the Intan headstage. An additional ground wire from the Faraday cage was connected to the ground wire of the Intan headstage to strengthen the ground connection. The probe was moved to the insertion position with the micromanipulator, guided by the microscope view. The actual probe insertion position was selected near the planned stereotaxic coordinates where no veins would be punctured. Prior to insertion, we measured the optical power entering the laser scanning system and subsequently performed the laser scanning calibration steps to ensure optimal light coupling into each core of the multicore fiber.

For experiments without the dura, probe insertion advanced at a speed of 1-2 *µ*m/s to minimize tissue damage [57] after confirming the probe was inserted. If the dura was left intact, during the first 500 - 900 *µ*m of insertion, the micromanipulator moved at a speed of approximately 0.5 - 1 mm/s to ensure a successful puncturing of the dura by the probe. If noticeable brain compression was observed under the microscope after dura puncturing, the probe was retracted back 200 - 500 *µ*m at a speed of around 10 *µ*m/s to relax the insertion pressure. Subsequently, we rested the probe in the brain for 10 - 20 minutes before further lowering the probe at an insertion speed of 1-2 *µ*m/s. Typically, the probe was inserted to a depth of 1 - 1.4 mm, resulting in the tip of the probe near or in the corpus callosum. After confirming that spontaneous and evoked spiking responses can be measured in the brain region, the probe rested in the position for 30 minutes to stabilize the signals prior to the photostimulation experiment.

After the experiment, the probe was retracted and immediately immersed in 1% Tergazyme solution (Sigma-Aldrich, St. Louis, MO, USA) for 2 hours. The probe was then rinsed with Milli-Q water for another 10-20 minutes. The power output from the probe was measured with a photodetector to monitor the coupling efficiency between the MCF and the chip. Additionally, the beam profile was visually inspected by projecting the beam onto a white card.

### I. Optogenetically induced seizure

The optogenetically induced seizure experiment was conducted with Thy1-ChR2-EYFP mice. All surgical and probe implantation procedures described in the previous section were applied, with two modifications. First, mice on P30 were injected with 10 nL of Kainic acid at 20 mM on the left side of the ventral hippocampus in the dentate gyrus (DG) region, following the same protocol described in [58] to develop an epilepsy mouse model. Second, the craniotomy was performed at the stereotaxic coordinates of AP: -1.6 mm and ML: 2 mm. The probe was inserted to a depth of around 1.7 mm to reach CA1 of the hippocampus. For the stimulation pattern, we used a 10 Hz pulse train with a 5-ms pulse width for 20 seconds, following the protocol from [50]. The stimulation pattern was repeated at various optical power levels starting from 19 *µ*W and increasing in steps of 16-21 *µ*W until the seizure was successfully induced. Mouse behavior after stimulation was recorded on video to score seizure severity.

### J. Electrophysiology recording configuration and photostimulation control

As the nanophotonic neural probe is passive, external instruments were required for the electrophysiology recording and control of the photostimulation pattern. To record electrophysiology signals, we used the Intan RHD 32-channel recording headstage for the LD probe, while a 64-channel version of the headstage (mini-amp-64, Cambridge NeuroTech) was used for the LS probe. Both headstages are compatible with the Open Ephys data acquisition board for saving and visualizing electrophysiology signals. The Open Ephys board was set to record wideband signals (1 - 7500 Hz) at a 30 kHz sampling rate per channel. Additional median subtraction and bandpass filter (300-6000Hz) were applied to the signal in the Open Ephys software for visualization of the spiking pattern in real-time. To control the photostimulation pattern, a custom MATLAB GUI was programmed to communicate with multiple instruments in the laser scanning system described in the Supplementary Section S2. Below is a list of the instruments used to control the photostimulation pattern:

- A MEMS mirror (A7B2.1-2000AL, Mirrorcle Technologies Inc., Richmond, California) in the optical scanning system directs light to the designated fiber core position on the MCF.
- A motorized rotational mount (K10CR1, Thorlabs) equipped with a variable neutral density (ND) filter (NDC-25C-2-A, Thorlabs), with an optical density (OD) range of 0.04-2, adjusts the input power to the scanning system.
- A microcontroller (Teensy 3.6, PJRC, Sherwood, OR) provides the TLL signal to modulate the laser diode (06MLD-488, Cobolt, Solna, Sweden) and the optical shutter (LS2S2T1, Uniblitz, Rochester, NY, USA). The microcontroller also sends the same TLL signal to the Open Ephys data acquisition board to label the stimulation time steps.
- A photodetector (S130C, Thorlabs) monitors the coupling efficiency of light to the fiber core on the MCF.

During the experiment, a set of stimulation patterns spanning various optical emitters and optical powers was defined in an Excel file. The program can read the Excel file to perform all the stimulation patterns in sequence. Throughout the experiment, the power entering the laser scanning unit was frequently monitored, followed by performing the feedback calibration, with intervals of less than 1 hour to ensure input power stability (see the Supplementary Section S2 for details).

### K. Electrophysiology data analysis

The wideband signals recorded from the Open Ephys board were post-processed in Python. First, the mean of the selected channel traces with minimal or no spikes was subtracted from all channels to reduce common noise interference. Subsequently, a third-order Butterworth bandpass filter with a passband of 300 Hz to 6000 Hz was applied to the signal. To account for the photovoltaic response induced by photostimulation due to various stimulation patterns (e.g., due to the pulse width, power, and channel), we separately calculated the average artifact waveform for each stimulation pattern and subtracted the artifacts induced by the same stimulation pattern, both at the onset and offset of the optical pulse. Furthermore, a blanking period of 1 ms before and 2.5 ms after the onset and offset of the optical pulses was implemented to ensure that the spike-sorting analysis would not detect these artifacts as spikes. An example of the processed traces is provided in Fig. S9b.

We performed spike sorting using the Spyking Circus package developed for sorting spikes for multichannel recording [59]. Subsequently, we performed a manual inspection of the sorted spikes with the phy GUI interface [60]. During the inspection, if unusual waveforms were detected in at least half of the channels for a particular unit, we identified it as an artifact and removed the corresponding time instances. The spike sorting algorithm was rerun to avoid compromising the clustering results due to the presence of artifacts. To prevent double counting of the same spike, we removed one of the paired spikes that have an interspike interval of *<* 0.5 ms. In the final results, all selected clusters met the following four criteria: 1) isolation distance *>* 10, 2) likelihood ratio *≤* 0.3 [27, 61], 3) SNR *≥* 3 [62], and 4) the percentage of spikes with refractory period violation (*<* 2 ms) *<* 2 % [57].

### L. Signal process on the LFP seizure responses

We applied common average referencing (CAR) to the raw signals with four of the channels that exhibited minimal seizure response as the reference signal, followed by bandpass filtering (5-300Hz). Calculation of the current source density (CSD) profiles was carried out following the procedure outlined in [63] with a modification of the spatial filter kernel size of the Hamming window set to 240 *µ*m (3 channels distance). Before performing the CSD calculation, we expanded the channel count from the original 6 selected channels to a total of 80 channels, each with a channel spacing of 5 *µ*m, with a channel interpolation method described in [64]. This interpolation enhances the resolution of the CSD heatmap.

### M. Histology verification of probe insertion position

To mark the insertion track of the probe in the tissue, the probe shank was coated with DiI red fluorescent dye (D-282, Thermo Fischer Scientific, Waltham, MA, USA) dissolved in ethanol (1-2 mg/ml) prior to insertion [65]. After the experiment, we extracted the mouse brain and prepared 300 *µ*m-thick coronal brain slices using a vibratome. To locate the position of the DiI insertion dye throughout the brain slices, an epifluorescence microscope equipped with an Alexa Fluor 568 filter cube and a YFP filter cube was used. The excitation wavelength was adjusted to 560 nm to visualize the DiI dye and 505 nm to visualize YFP expression in brain tissue. YFP images of the brain were overlaid with DiI fluorescent images to verify the probe insertion position. We also captured images of the entire coronal brain slices to facilitate alignment with the corresponding coronal section in the mouse brain atlas.

### N. Statistical analysis

The statistical analysis was conducted using either Python or MATLAB. All statistical comparisons between the groups were performed with the two-tailed non-parametric Mann-Whitney test unless otherwise stated. Results were reported with statistical significance if they met the threshold of p *<* 0.05.

## V. DATA AVAILABILITY

All simulation data, raw and processed electrophysiology data reported in this study are available at https://doi.org/10.17617/3.ZX0YAE. Additional data are available upon reasonable request from the corresponding authors for scientific use.

## VI. CODE AVAILABILITY

The Python codes for electrophysiology data analysis and the MATLAB codes, mainly for plotting, are posted at https://doi.org/10.17617/3.ZX0YAE. Additional codes are available upon reasonable request from the corresponding authors for scientific use.

## ACKNOWLEDGMENTS

F.D.C. thanks M. Brunk for sharing the Matlab script used for CSD calculation, A. Stalmashonak and F. Weiss for assistance with the optical measurements and MCF-to-chip attachment setup, P. Shah (Krembil Research Institute) for his help in the initial *in vivo* experiments, H. Steenland (NeuroTek) and Y. Chen (Max Planck Institute for Biology) for suggestions on the animal experiments, and M. Lippert and E. Duŕan (Leibniz-Institut für Neurobiologie, Magdeburg) for discussions on spike sorting. J.K.S.P. thanks M. Roukes (Caltech) for fruitful discussions and J. Siegle (Allen Institute) for input on electrode impedance reduction. Funding support from the Max Planck Society, Natural Sciences and Engineering Research Council of Canada, and the Canadian Institute of Health Research is gratefully acknowledged.

## CONTRIBUTIONS

W.D.S. and J.K.S.P. conceived the initial idea. F.D.C. and T.X. performed the device simulations, designed the probes with input from W.D.S. and J.K.S.P., and laid out the designs. H.C., X.L., and P.G.Q.L. were responsible for the wafer fabrication. H.W. designed and constructed the MEMS-based laser scanning system. F.D.C. characterized the devices with the help of Y.J. in investigating the electrode structure. W.D.S. proposed the laser treatment for electrode impedance reduction, and the method was developed by F.D.C.. F.D.C. performed the device packaging with support from J.N.S. for the wirebonding and X.M. and P.D. for the fiber-to-chip attachment process. W.D.S, A.S., and A.G. developed the polishing process for thinning the probe. F.D.C., H.M.C., M.M., and D.A.R conducted the animal experiments. F.D.C. analyzed the data with guidance from H.M.C. and J.K.S.P.. F.D.C. and J.K.S.P. co-wrote the manuscript with inputs from other co-authors. The project was supervised by T.V., W.D.S., and J.K.S.P..

## SUPPLEMENTARY INFORMATION

### Supplementary Video 1

Side-view fluorescence microscope imaging of an LD probe immersed in a fluorescein solution. We switched between all 16 emitters on the probe using the MEM-based laser scanning system. The power ratio between the emitters with the highest and the low-est output power was 2.7 times. The video is in real-time. The video is available at: https://doi.org/10.17617/3.ZX0YAE.

### S1. SHANK THICKNESS WITH POST-PROCESS BACK POLISHING

As described in Methods of the main text, in addition to the backgrinding performed on a wafer scale at the foundry to thin the wafer to *∼* 100 *µ*m, we performed a second polishing process on a singulated probe to further reduce the probe thickness. Figure S1a shows the side-view micrograph of a probe shank before and after the second polishing step. To assess the effectiveness of this process, we compared the shank thickness of five unpolished probes with that of five polished probes in Figure S1b. All probes included in the box plots are from the same wafer. The thicknesses of the unpolished probes were 98 - 105 *µ*m while the thicknesses of the polished probes ranged from 37 to 62 *µ*m, achieving on average *∼* 2*×* thickness reduction.

**FIG. S1.**
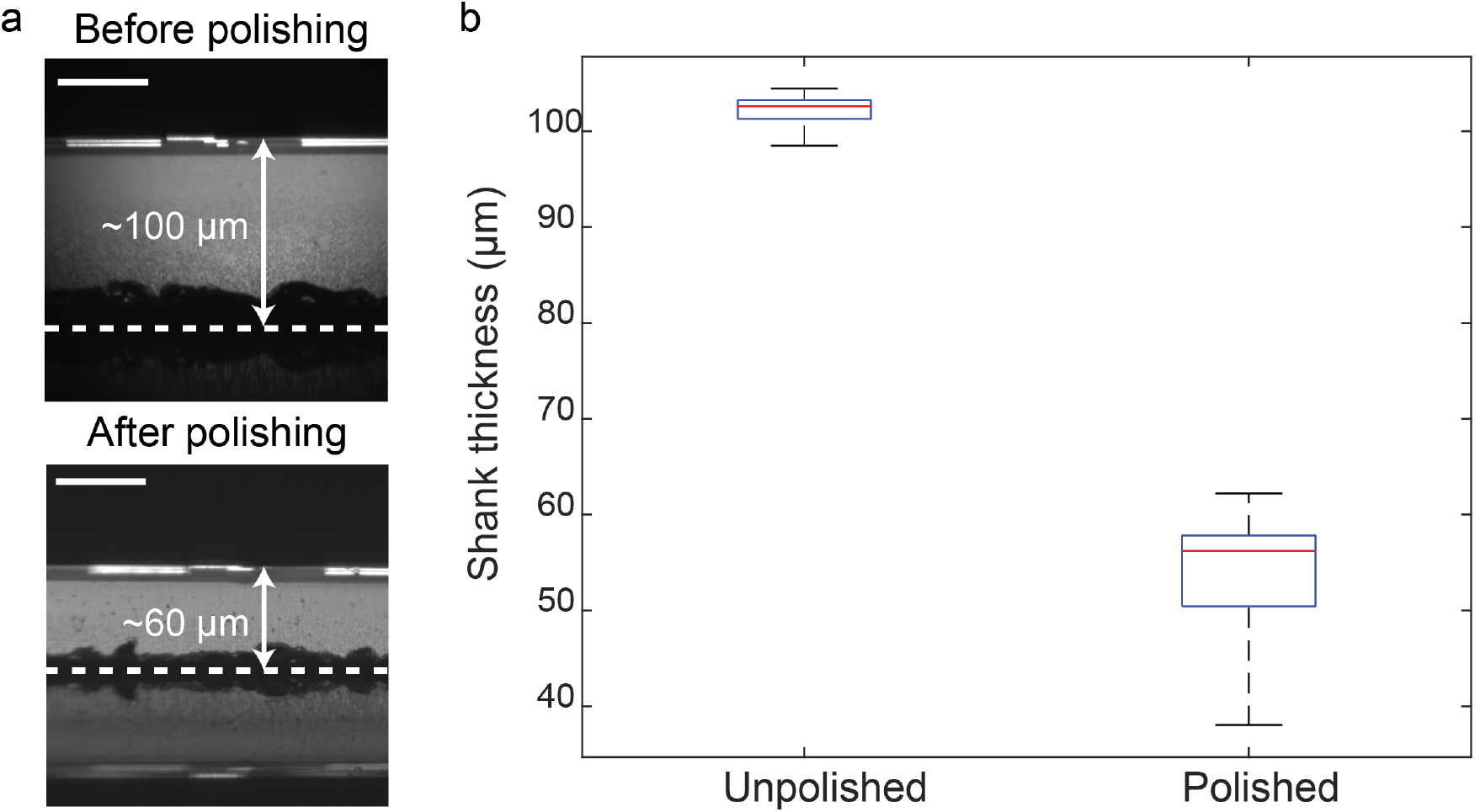
Comparison of the probe thickness before and after the polishing process in a fiber polisher. **a** Side view micrographs of an LD probe before and after the backgrinding process, showing a reduction in probe thickness from 104 *µ*m to 58 *µ*m. The scale bars are 50 *µ*m. The white dashed lines indicate the bottom surface of the probe estimated based on the center position between the probe and its bottom reflection image. **b** Thickness comparison of 5 probes without and another 5 probes with the additional polishing process in box plots, with red lines representing the median, box boundaries representing the interquartile range, and the whisker representing the minimum and maximum thickness.

### S2. LASER SCANNING UNIT

Figure S2a shows a schematic of the laser scanning unit used to couple light into the probes in the experiments. This setup is an extension of our previous laser scanning unit reported in [S2, S3]. Here, we have implemented three additional features to improve the power stability of coupling light into the MCF for animal experiments that last hours. First, we reduced the length of the scanning system from *≈* 60 cm to 10 cm with custom lenses to improve the mechanical stability, as shown in Fig. S2b. Second, we made the scanning system less susceptible to polarization drift, primarily due to the extended length of the MCF (exceeding 1 m). We achieved this by depolarizing the input light using the following method: the half-wave plate (HWP) with the polarization beam splitter (PBS) splits the input light into two orthogonally polarized beams with equal power. By ensuring that the path length difference between these two beams exceeded the laser coherence length, the recombined beam effectively became depolarized. Third, we introduced a feedback system to maintain optimal coupling efficiency between the scanner and the MCF. By minimizing scattered light in the MCF cladding, we effectively maximize the efficiency of coupling light to the fiber core, as demonstrated in Figure S2c. A gradient descent-based algorithm was developed to control the MEMS mirror coordinates to minimize the scattered light. The algorithm takes around 1 - 2 s to optimize the MEMS coordinates for one channel, and we manually triggered the feedback algorithm at an interval of less than 1 hour during the experiment to ensure consistent output power from the probe. We expected the drift of the output power of the probe to be less than 10 % in 1 hour, as shown in Fig. S2d. During the experiment, to prevent the light used in the feedback system from stimulating the brain, we reduced the input power by introducing an additional ND filter (OD 1) and setting the variable ND filter to OD 1.5-2. In future animal experiments, this same principle could be applied to develop a closed-loop feedback system capable of continuously tracking and maintaining the coupling efficiency, as demonstrated in Fig. S2e.

**FIG. S2.**
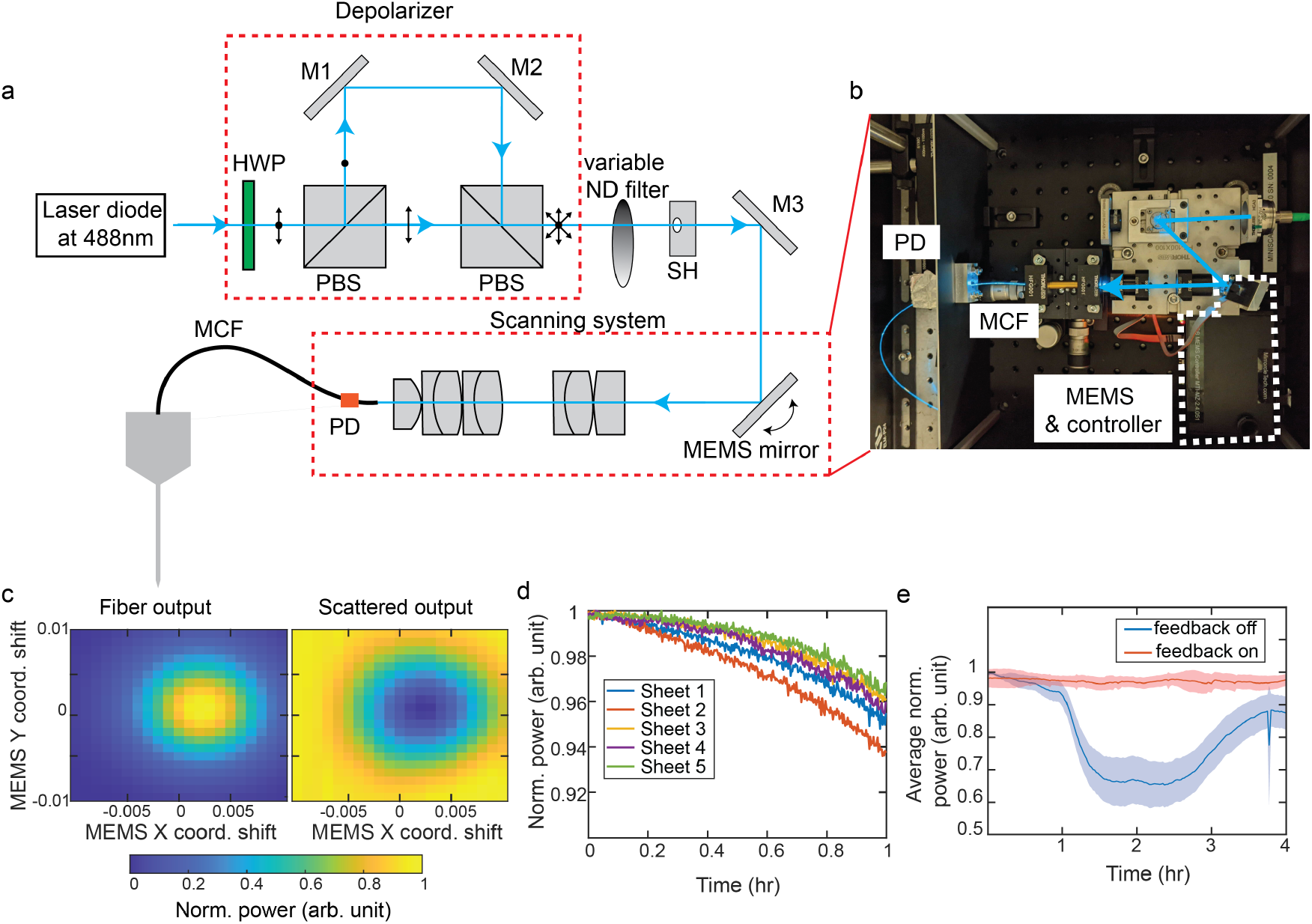
The MEMS-based scanning system for addressing different fiber cores on MCF. **a** Schematic of the optical scanning unit with the depolarizer (not drawn to scale). HWP: half-wave plate, PBS: polarization beam splitter, M: optical mirror, SH: optical shutter, PD: photodetector. **b** Image of the compact scanning system. For the animal experiments, the depolarizer was implemented outside of the scanning system, and they are connected through a fiber. However, they can be combined into a single free-space optical system, as demonstrated in our previous work in [S1]. **c** Comparison of MCF output power and optical power scattered in the fiber cladding immediately after MCF input for various MEMS position coordinates (coord.). We optimized the light coupling efficiency to each fiber core by minimizing the scattering power. **d** Average output power of individual light sheet emitters on an LS probe over 1 hour, averaged over 7 repetitions. On average, the power dropped by less than 10 % across all 5 emitters on the LS probe. **e** Average normalized output power of all 16 emitters on the LD probe over a 4-hour period with and without the closed-loop feedback mechanism that utilizes light scattered in the fiber cladding to optimize light coupling to the MCF cores. The shaded region indicates the SD of the output power.

### S3. ELECTRODE IMPEDANCE REDUCTION WITH DIFFERENT SOLUTION TREATMENTS

To investigate the effect of electrode impedance reduction following immersion in 1% Tergazyme(pH = 9.3), four LD probes with 20 *×* 20 *µ*m^2^ electrodes were immersed in three different solutions for a minimum of 5 hours. These solutions included 1% Tergazyme, water with a pH of 9.5, and 1*×* PBS. Laser treatment was applied to the electrodes on three probes, leaving one untreated for comparison. Figure S3a illustrates the comparison of electrode impedance before and after different solution treatments applied to each probe. Here, we observed similar impedance reduction between the probes with laser treatment and soaked in Tergazyme and in basic water. This result suggests that the impedance reduction associated with Tergazyme washing may be attributed to its basic nature rather than the presence of protease enzymes. We also observed that the laser-treated electrodes experienced a decrease in impedance after soaking in PBS for 8 hours, though the effect was less pronounced than in the basic solutions. Lastly, electrodes without laser treatment also had an impedance reduction after Tergazyme soaking, but Tergazyme alone did not reduce the electrode impedance below the 2 MΩ threshold, confirming the necessity of our electrodes to be laser treated. Further structural analysis, conducted using both an optical microscope and scanning electron microscope, on the electrodes before and after Tergazyme immersion, presented in Figs. S3b and c, reveals no noticeable surface degradation. The actual mechanism of the impedance reduction with Tergazyme treatment is yet to be determined.

**FIG. S3.**
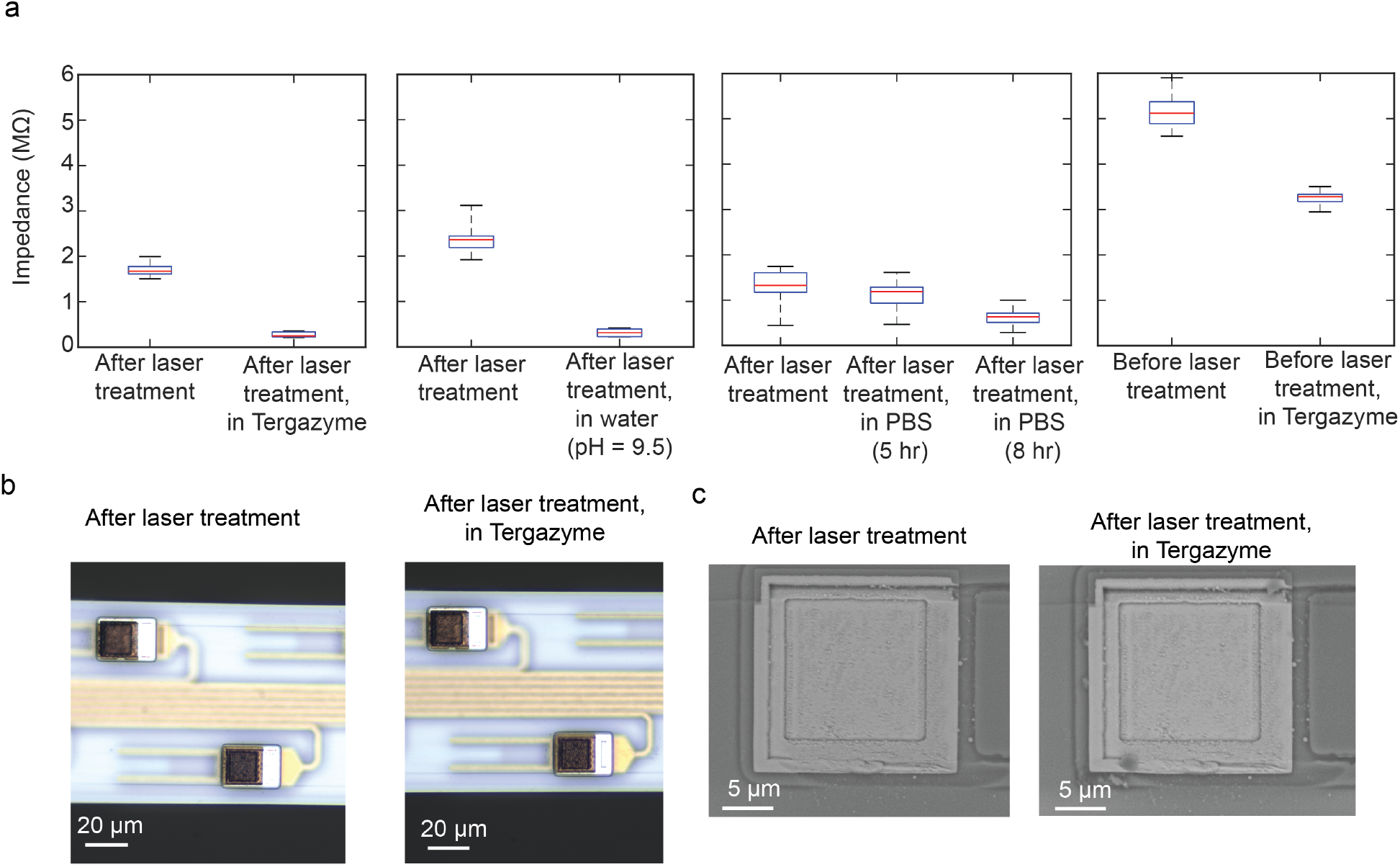
**a** Comparison of the electrode impedance for 4 LD probes that underwent different solution treatments. The probes were completely immersed in the specified solution for 5 hours, unless otherwise specified, before measuring the impedance in PBS. The data are presented in box plots with the box indicating the interquartile range (IQR) and the red line denoting the median impedance. Whiskers extend to the minimum and maximum electrode impedance on the probe. **b** Optical micrographs of the electrodes on an LD probe before and after soaking in Tergazyme solution for 5 hours. **c** SEMs of an electrode on a different LD probe before and after soaking in Tergazyme for 5 hours. The scale bars are 20 *µ*m in **b** and 5 *µ*m in **c**.

### S4. OPTICAL CHARACTERIZATION OF THE SYSTEM

Figure S4 summarizes the insertion loss of the grating emitters in the probes used in the animal experiments. The transmission of the emitters varied between -34 and -19 dB across the probes, while the transmission variation between the emitters on each probe ranged from 0.2 to 5.8 dB. In particular, this power distribution is significantly more uniform compared to using an imaging fiber bundle with irregular fiber core positions for coupling light to the chip [S4] (maximum variation *∼* 16dB).

**FIG. S4.**
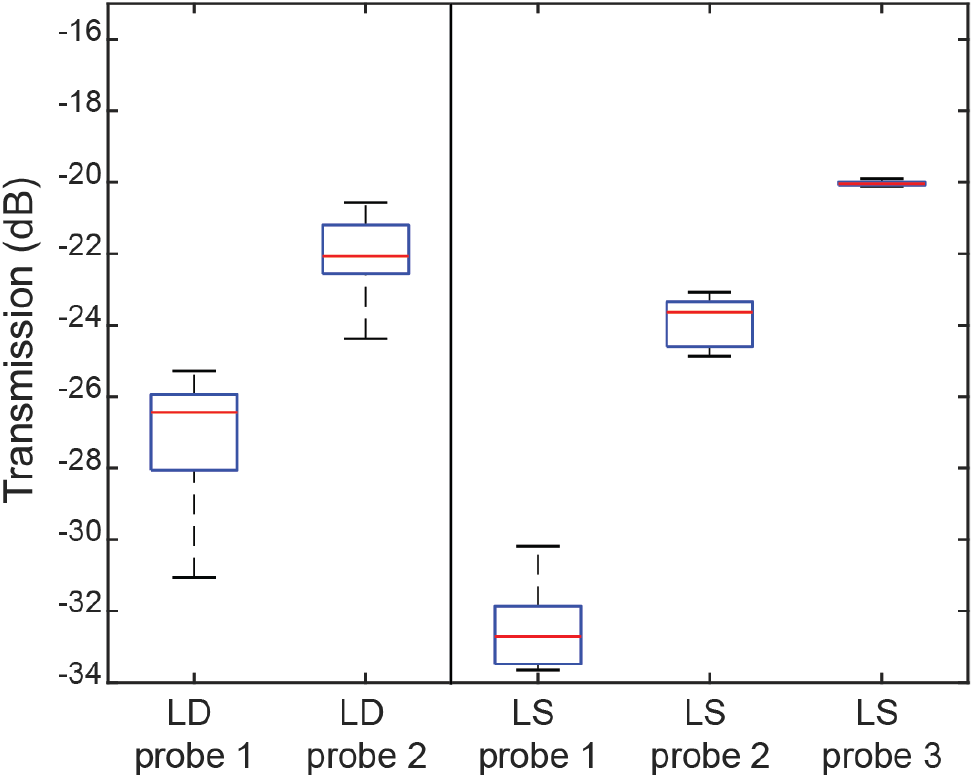
Box plots of the optical transmission of the packaged probes used in the *in vivo* experiments. Input power measurements were taken at the fiber input into the scanning system, while output power measurements were recorded at the emitter. Each box plot includes all emitters on each probe (LD probe, n = 16 emitter; LS probe, n =5 emitters), except for two emitters on LD probe 1, which were excluded due to damaged edge couplers. The box represents the interquartile range (IQR) of optical transmission for the emitters on the probe, with the red line denoting the median value. Whiskers extend to the minimum and maximum optical transmission values recorded for the emitters on the probe

The insertion loss of the neural probe is primarily attributed to the low edge coupling efficiency (7-11 dB [S2]) and the MCF-to-chip alignment drift during and after packaging, partly because the edge coupler had a large mode mismatch with the fiber and a tight alignment requirement (3dB tolerance of *<* 2 *µ*m), as investigated in [S5]. The LS probe suffered an additional *∼* 5dB loss compared to the LD probe due to the on-chip waveguide routing network, which distributed an edge-coupled input to 8 grating emitters on 4 shanks.

Despite the high optical loss observed in LD probe 1 and LS probe 1, both probes could still provide adequate optical intensity to elicit spiking responses. Our external laser scanning system could deliver at least 15 mW with short pulses (*≤* 30 ms) without an immediate decrease in the output power of the probe. This translated to a minimum output of 6 *µ*W for the worst emitter on LS probe 1. However, it should be noted that we observed a sudden decrease in optical transmission at high input powers (*>* 60 mW with a 30 ms pulse width), potentially attributed to the degradation of the epoxy used for MCF-to-chip attachment, as also mentioned in [S6].

The optical insertion loss of the neural probe can be improved by employing more efficient bi-layer edge couplers [S5], and optimizing the MCF-to-chip attachment process (i.e., improving mechanical stability of the fiber and probe clamps, testing different epoxies, etc.). As is evident in Figure S4 where probes with larger index numbers were packaged at a later time, the transmission of the LS probe improved by an order of magnitude simply by fine-tuning the MCF-to-chip attachment procedure. We expect the LD probe to reach an optical insertion loss of *<* 13 dB with these two improvements. A systematic analysis of the packaging yield can be carried out in future work. Also, MCF-to-chip attachment using a semi-automatic machine described in [S1] can help improve the packaging throughput and repeatability.

**FIG. S5.**
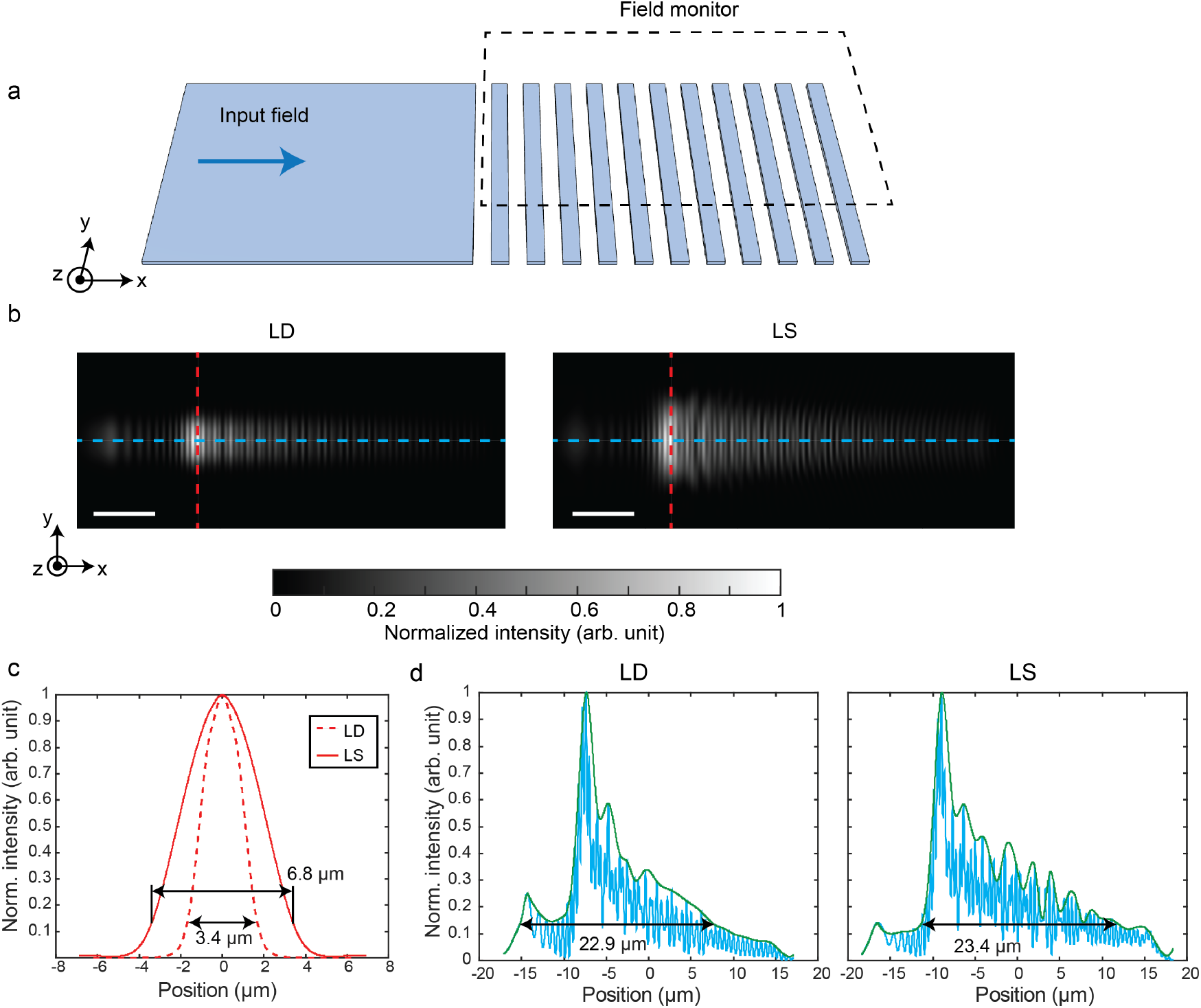
**a** Illustration of the setup to simulate the beam emission profile from the grating emitter. **b** The near field beam emitted by a low-divergence emitter and one of the eight light sheet emitters. To simulate the depolarized near-field pattern from the emitter, we superimposed the TE and TM polarization emission patterns with their intensities adjusted based on the output coupling measurements for each polarization. The scale bars are 5 *µ*m. **c** Normalized intensity along the vertical line cuts (red dashed lines in **b**) of the low-divergence and light sheet beams. **d** Normalized intensity along the horizontal line cuts (blue dashed lines in **b**) of the low-divergence and the light sheet beams. The green lines represent the envelopes used to estimate the beam widths. The size of the low divergence and the light sheet beams measured at 1/e^2^ of the peak intensity are 22.9 *×* 3.4 *µ*m^2^ and 23.4 *×* 6.8 *µ*m^2^, respectively.

### S5. ESTIMATION OF BEAM INTENSITY PROFILE

We estimated the beam intensity distribution of the two beam types at a specified output power in units of mW/mm^2^ using the simulated and measured normalized beam intensity profiles in cortical slices. For the simulation estimation, we used the BPM described in Methods to acquire the volumetric beam profile with scattering properties similar to those of the cortex. To convert the normalized intensity value to mW/mm^2^, we measured the beam size, as indicated by the 1*/e*^2^ width and length of the input beam, as shown in Figs. S5b-d. Subsequently, we computed the average intensity within the beam. The ratio of this average intensity value to the actual output beam intensity calibrated the intensity distribution of the entire beam profile. The top-down intensity profile, as shown in Fig. S6a, was extracted from a transverse plane at the depth with the highest intensity at the input to the scattering medium.

**FIG. S6.**
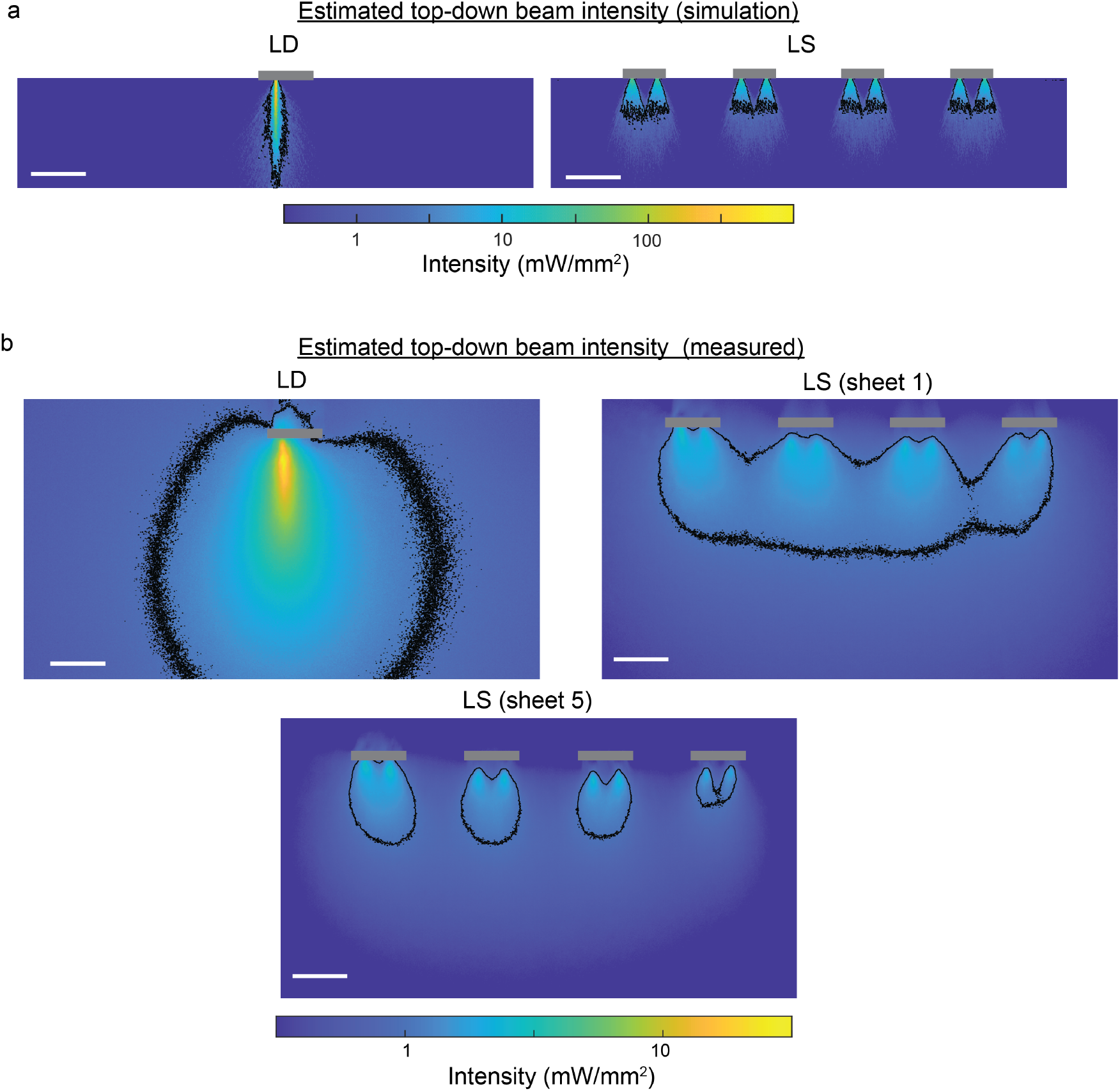
Estimated top-down beam intensity profile at an output power of 10 *µ*W calculated with **a** simulated beam profile in scattering medium and **b** measured beam profile in cortical brain slices. Sheet 5 beam profile in **b** has a non-uniform beam intensity between the right-most and left-most emitters because of the additional waveguide crossings to route light to the right side of the probe. The maximum power variation between the emitters is 2*×*. The images are plotted in log scale, and the black contour lines label the 1 mW/mm^2^ intensity level. The grey boxes show the positions of the probe shanks. All scale bars are 100 *µ*m.

To convert the experimental beam profile intensity captured with the epifluorescence microscope to absolute intensities (with units of mW/mm^2^), we first measured the beam size at the emitter output in the beam profile images in tissue. We estimated the beam width with 1*/e* of the peak intensity instead of 1*/e*^2^ because of the higher background signal measured in the tissue image. The beam thickness was approximated using the side-view beam profile measured in the fluorescein solution, as shown in Fig. S7. Considering that the side-view profile of a light sheet beam overestimates the beam thickness caused by out-of-focus light, we approximated the sheet thickness using the low-divergence beam profile. This approximation was valid, as the two beam types exhibited similar beam thicknesses, as shown in the simulations in Figs. S5b and d. Finally, we calculated the average intensity along a line profile at the peak intensity point near the emitter, with the line length matching the measured beam width. With the estimated beam size and the average intensity value at the output beam emitter, we were able to reference the normalized intensity to the absolute intensity in mW/mm^2^ at a given output power.

The estimated beam intensity profiles for both beam types at an output of 10 *µ*W are displayed in Fig. S6. A notable difference between the simulated and experimental beam intensity profiles is the peak intensity value. The lower intensity value in the experimental results can be attributed to an overestimated beam width due to out-of-focus light and additional scattering caused by the beam at depth in tissue. The fluorescein-stained fixed tissue also leaked some fluorescein into the PBS solution, further increasing the background signal. This larger background signal also enlarged the effective stimulation area highlighted with the 1 mW/mm^2^ contour line. Nevertheless, the light sheet beam consistently exhibited lower output intensity compared to the low-divergence beam of the same estimation approach at the same output power. The difference in the peak output intensity between the two beam profiles was 15.7*×* in simulation (LD: 580.1 mW/mm^2^, LS: 36.9 mW/mm^2^) and 9.7*×* in experimental results (LD: 28.1 mW/mm^2^, LS: 2.9 mW/mm^2^).

**FIG. S7.**
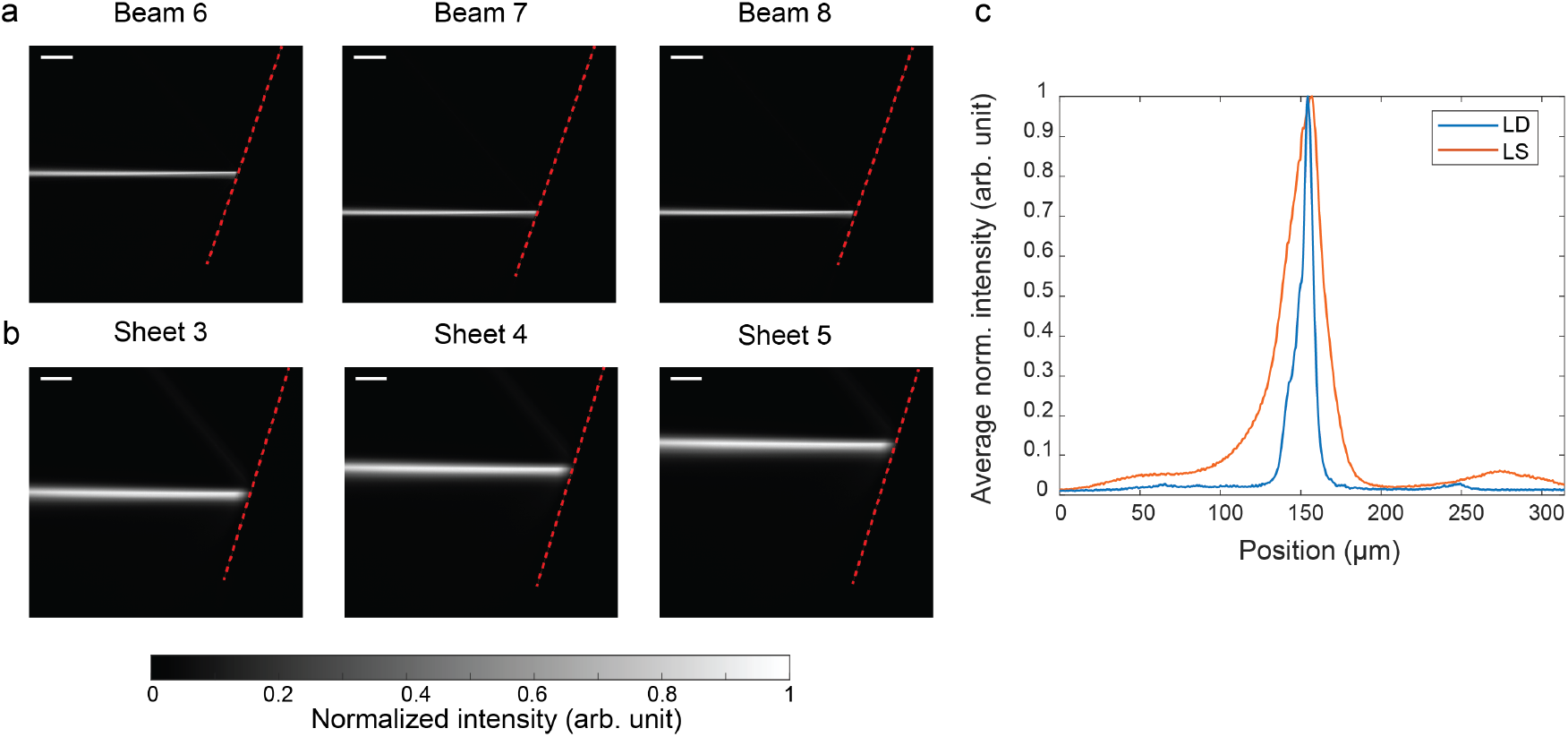
Side view beam profile of **a** the low-divergence beam and **b** the light sheet beam measured in a 10 *µ*M fluorescein solution. Three different emitters on the probes were addressed with no visible optical crosstalk between channels. The red dashed line indicates the surface of the probe. The scale bars are 100 *µ*m. **c** An average normalized line profile from five emitters of each probe type, measured at a distance of 100 *µ*m from the emitter. The FWHM for the low-divergence beam and the light sheet beam are 9.4 *µ*m and 26.3 *µ*m, respectively.

### S6. SIDE VIEW BEAM PROFILES IN FLUORESCEIN SOLUTION

To investigate the beam thickness of the low-divergence beam and the light sheet beam, we immersed the probes in fluorescein solution and captured a side view using a fluorescence microscope with a green fluorescent protein (GFP) optical filter. Figures S7a and b show the side view beam profiles for three emitters on the LD and LS probes. An average background value estimated with an image taken in the absence of beam emission was subtracted from the presented side-view images. Overall, only a single forward propagating beam was visible from each emitter, demonstrating minimal crosstalk between channels. The peak intensity of the second-order diffraction beam from the grating emitter was at least 10 dB lower than that of the main beam. The line beam profile measured at a propagation distance of 100 *µ*m shows the light sheet beam was *>* 2 *×* thicker than the low-divergence beam. However, the measured sheet thickness based on the side-view beam profile is an overestimation due to the stronger out-of-focus light stemming from the wider extent of the sheet width. From our previous sheet thickness measurement by projecting the sheet onto a coverslip coated with a fluorescent thin film in air [S4], the sheet thickness is expected to be 8 to 12 *µ*m, similar to the low-divergence beam thickness.

### S7. HISTOLOGICAL VERIFICATION OF IMPLANTED LOCATIONS

Figure S8 shows 3 histology images, each from an animal experiment, with the probe insertion track labeled with the DiI fluorescent marker overlaid on top of the corresponding coronal brain YFP image. Probe positions were first estimated with the whole coronal brain slice camera images to identify the position of the cross-sectional plane. Subsequently, fluorescence histology images were used to precisely determine the location of the insertion track relative to the distinctive structures nearby. The histology images in Figures S8a and b confirmed that the LS and LD probe experiments discussed in Sections II C and II D of the main manuscript were inserted into layers V and VI between the motor cortex and the somatosensory cortex. In Figure S8 c, the histological image verified the seizure induction in CA1 of the hippocampus discussed in Section II E of the main text.

### S8. INVESTIGATION OF LIGHT-INDUCED ARTIFACTS AND SIGNAL PROCESSING FOR ARTIFACT REMOVAL

We investigated the photostimulation response in a control experiment with an LD probe implanted in an anesthetized wild-type mouse, targeting the same brain region as described in Sections II C and II D of the main manuscript, located between layers V and VI of the motor and somatosensory cortex. In Figure S9a, it is evident that no spikes were observed during the optical pulse in the absence of opsins. The notable large spikes that appeared at the onset and offset of the optical pulse are attributed to the photovoltaic effect. The amplitude of these artifact spikes varied according to the distance of the electrode to the activated grating emitter, as shown in Fig. S9b. Electrodes closer to the stimulation emitter experience a stronger photovoltaic effect, resulting in larger artifact amplitudes.

To ensure accurate spike detection without miscounting spikes, we applied an average artifact subtraction procedure on each channel. This method served to minimize both the amplitude and duration of the artifacts. The average artifact was calculated using artifacts generated by optical pulses with the same power and duration. Additionally, we implemented a blanking period of 1 ms before and 2.5 ms after the optical pulse to effectively mask the presence of light-induced artifacts. With these artifact removal steps in place, we were able to retain most of the spiking activity while minimizing the impact of the light-induced artifact on spike detection, as illustrated in the example voltage traces in Figure S9.

**FIG. S8.**
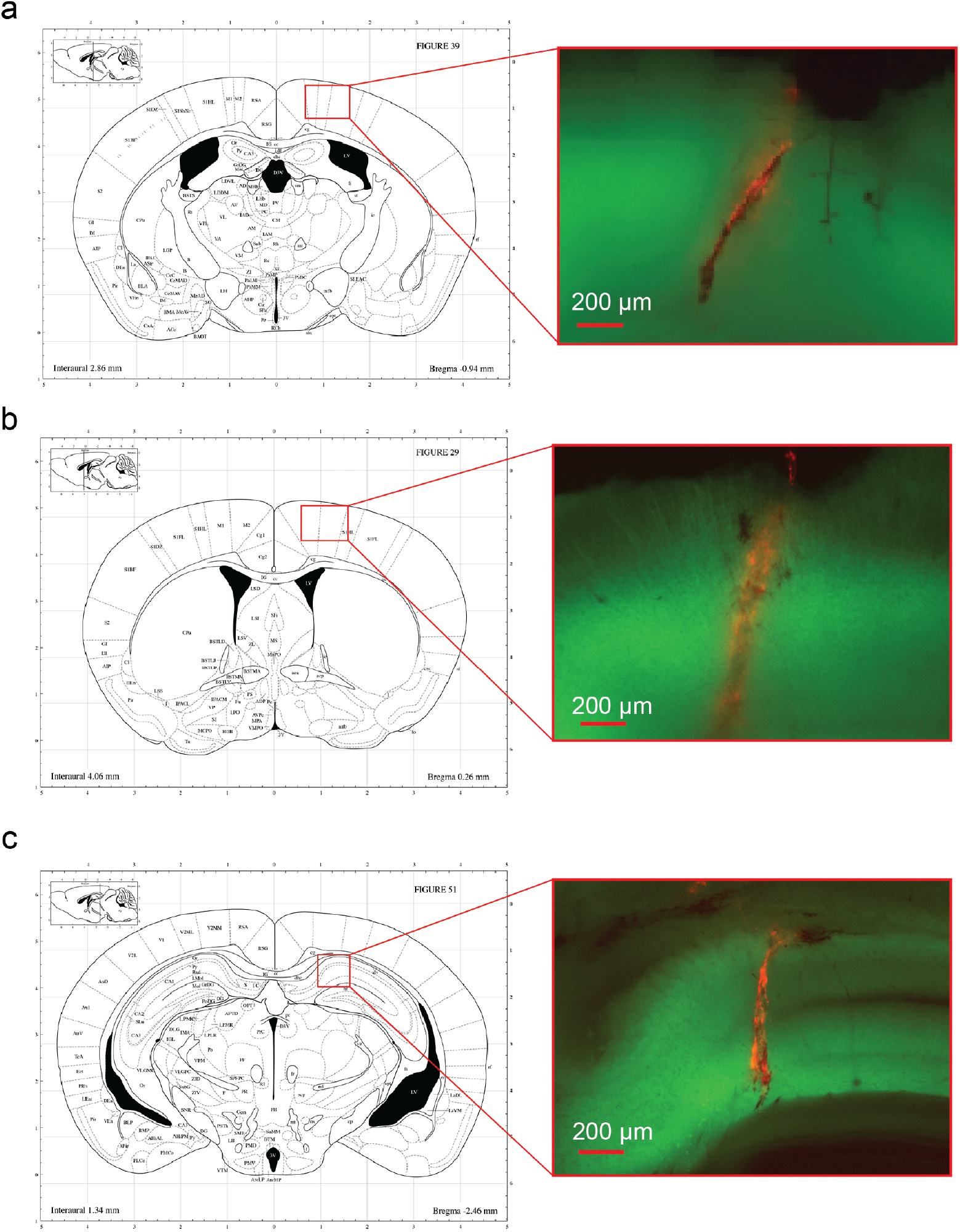
Cross-sectional 2D representations of the mouse brain, illustrating the estimated probe implantation locations for three animal experiments: **a** the LS probe experiment in Fig. 4a and 5a, **b** the LD probe experiment in Fig. S10, and **c** the optogenetically induced seizure experiment in Fig. 6. The insets show YFP fluorescence images of coronal brain slices (in green) overlaid with probe insertion tracks marked using DiI fluorescent dye (in red). The 2D drawings of the coronal mouse brain section are reproduced from [S7].

**FIG. S9.**
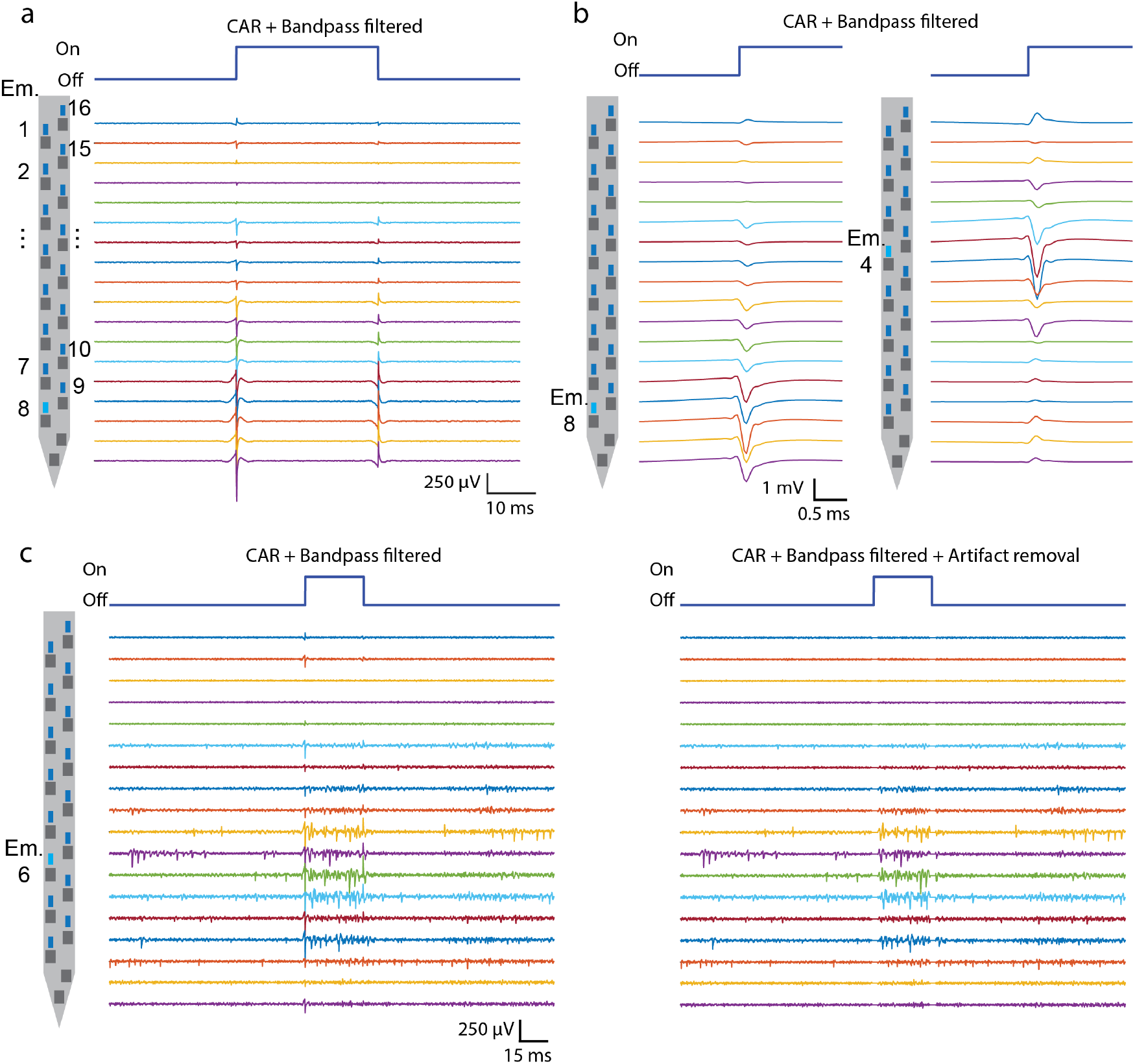
**a** Snapshot of filtered voltage traces (common averaging remove (CAR) + 300 - 6000 Hz) recorded with an LD probe in an anesthetized wild-type mouse. A 30 ms optical pulse at 4.9 *µ*W was applied to Emitter (Em.) 8 labeled in light blue. No spiking response was observed during the optical pulse. The two large spikes at the onset and offset of the optical pulse were caused by the photovoltaic effect. **b** Average light-induced artifacts caused by 2 emitters. The voltage traces were averaged over 30 optical pulses. The optical power applied to Em. 8 and Em. 4 were 4.9 *µ*W and 6.1 *µ*W, respectively. **c** Snapshot of voltage traces recorded in the cortex of a Thy1-ChR2 mouse before and after removal of the light-induced artifact. The artifact removal procedure includes artifact subtraction and blanking for a short period at the onset and offset of the stimulation pulse.

### S9. SPATIALLY SELECTIVE STIMULATION WITH AN LD PROBE

Similar to the analysis conducted in Figure 4b - e, Figure S10 shows spatially selective stimulation using an LD probe in an awake head-fixed animal experiment. The LD probe was inserted between the motor and somatosensory cortex of a Thy1-ChR2 mouse. Most electrodes were placed within layers V and VI of the cortex, as visualized in Figure S10a. The electrodes in layer II/III and above tended to record low-amplitude spikes or no spikes. Figure S10b shows the photostimulated spiking responses elicited by Em. 7 and 3 on the probe. The spiking response shifted towards the electrodes at a shallower depth when the light was switched from Em. 7 to Em. 3. Additionally, Figure S10c confirms that neurons exhibiting a high ActProb (*>* 90%) were in close proximity to the addressed emitters.

When examining the two example units presented in Figure S10d and e, we found that the neuron firing rate during the stimulation optical pulse can return to the baseline firing rate after stimulating with an emitter that was 3-5 emitters away (384 - 640 *µ*m) from the location that evoked the highest spike rate. The longer stimulation distance observed in this particular example compared to Section II C of the main text could be attributed to the higher output power of each LD emitter (LS: 2.1 - 2.5 *µ*W, LD: 3 - 4 *µ*W), or it may have resulted from the higher output intensity of the LD emitter, which could lead to longer beam propagation distances. In general, this experiment demonstrates the capability of an LD probe to selectively stimulate different brain regions along the probe shank. When examining the two example units presented in Figure S10d and e, we found that the neuron firing rate during the stimulation optical pulse can return to the baseline firing rate after stimulating with an emitter that is 384 - 640 *µ*m (equivalent to 3-5 emitters) away from the location that evoked the highest spike rate. The longer stimulation distance observed in this particular example compared to Section II C of the main text could be attributed to the higher output power of each LD emitter (LS: 2.1 - 2.5 *µ*W, LD: 3 - 4 *µ*W), or it may have resulted from the higher output intensity of the LD emitter, which could lead to longer beam propagation distances. In general, this experiment demonstrates the capability of an LD probe to selectively stimulate different brain regions along the probe shank.

**FIG. S10.**
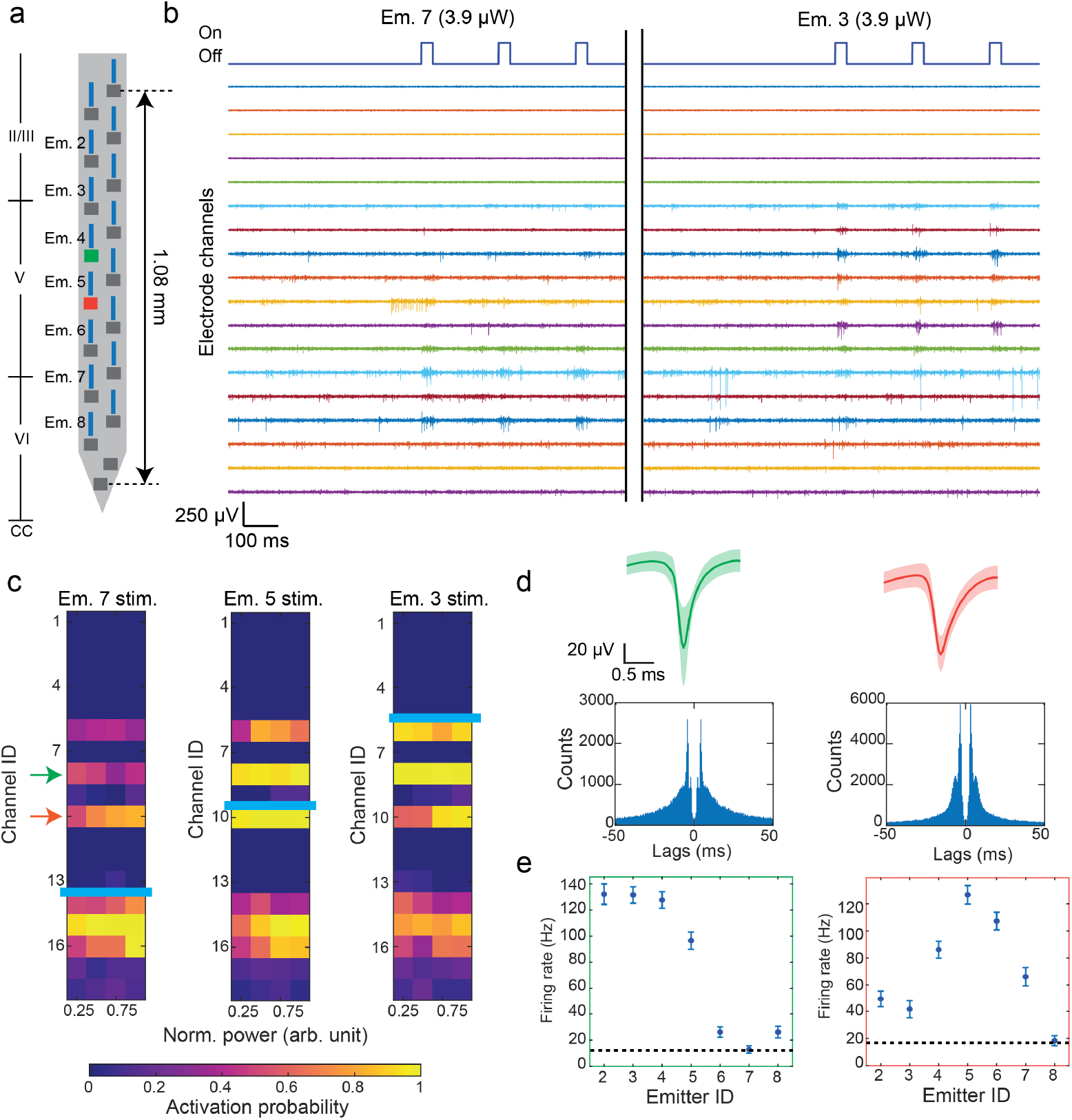
*In vivo* demonstration of selective optogenetic stimulation using the LD probe in an awake head-fixed mouse. **a** Estimated implantation position of the LD probe within the cortex of a Thy1-ChR2 mouse, with labels indicating the indices of the emitter channels used in the experiment. **b** Snapshot of the example voltage traces from all 18 electrode channels on the probe during a 30 ms pulse train photostimulation from two of the emitters. Em. 7 and 3 induced spiking activities in channels at different depths. **c** Mean ActProb of neurons recorded by the LD probe with increasing photostimulation powers of Em. 7, 5, and 3. Power levels are normalized to the trial with maximum output power (Em. 7: 5.3 *µ*W, Em. 5: 7.6 *µ*W, Em. 3: 8 *µ*W). The position of the stimulation emitter is indicated by a light blue line. Neurons with an ActProb *>* 90% are close to the stimulation emitter.**d** Mean waveforms and autocorrelograms of two example units recorded on the LD probe. The shaded area superimposed on the waveforms represents the SD of the waveform. The colors correspond to the arrows in **c**, which denote the positions of the neurons. **e** The mean firing rate of the two example units evoked by different emitters. The output power of each emitter ranged between 3-4 *µ*W. The black dashed lines indicate the average firing rate at baseline, calculated at 10 s before each stimulation pulse train. The error bars denote s.e.m.

**FIG. S11.**
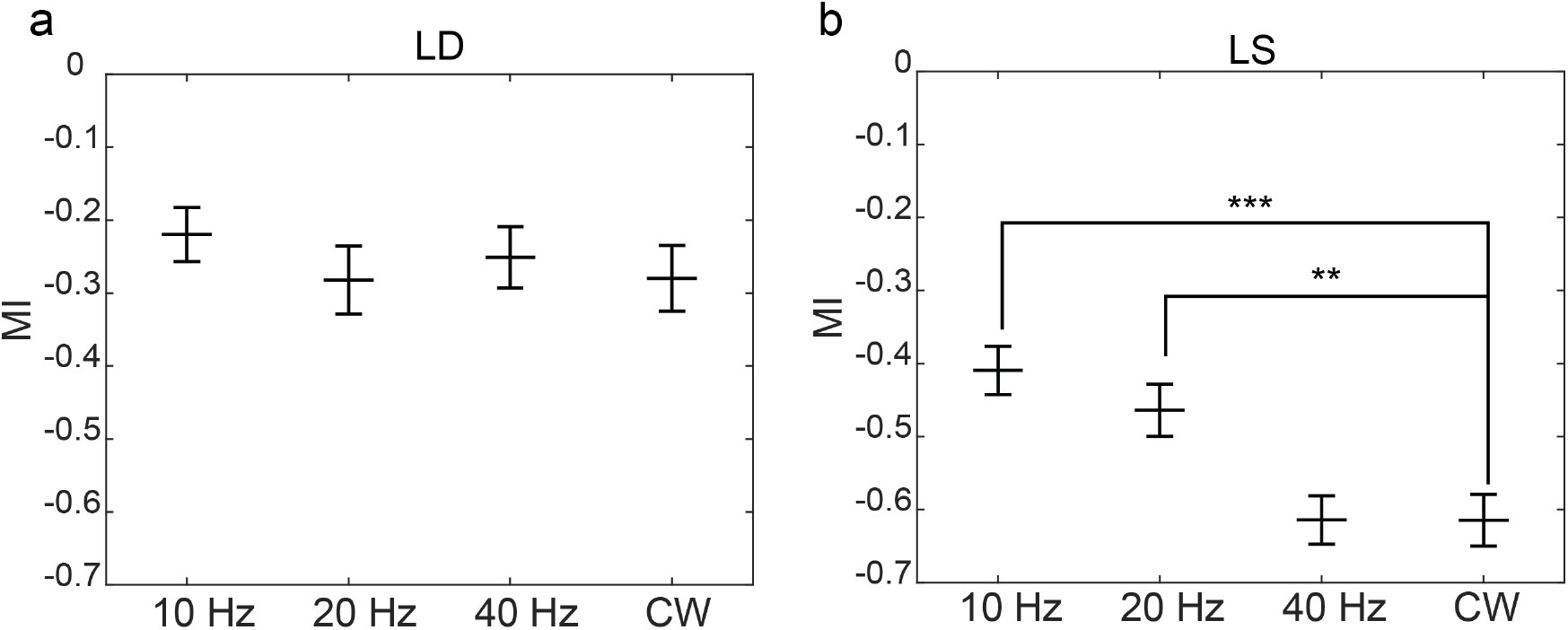
Comparison of the MI of the identically selected stimulated units presented in Fig. 5d and e elicited by **a** LD and **b** LS probes for different stimulation patterns. 10 ms-long optical pulses were applied at frequencies of 10, 20, and 40 Hz for 1 s. CW represents a continuous 1 s-long stimulation pattern. Each stimulation pattern was repeated 10 times, with 20 - 30 s intervals between repetitions. One-tailed non-parametric Mann–Whitney test was employed to statistically compare the inhibition responses induced by CW stimulation with the responses induced by stimulation at other frequencies. ** denotes p = 0.0011 and *** denotes p *<* 0.001.

### S10. COMPARISON OF THE INHIBITION STRENGTH AFTER PHOTOSTIMULATION

In addition to investigating inhibition responses following the 1 s continuous wave (CW) stimulation pattern, we studied the spiking responses modulated with 10 ms optical pulse trains at frequencies of 10, 20, and 40 Hz, each lasting 1 s. Each of these stimulation patterns was repeated 10 times with an interval of 20-30 s during the same animal trials (LD: 6 animals, LS: 7 animals). Figure S11 presents the statistical comparison of inhibition strength after different stimulation patterns quantified using MI. The same selected neurons reported in Figures 5 d and e are included in this comparison. In particular, we observed that stimulation patterns at 40 Hz and CW can lead to a lower mean modulation index (MI), with a greater statistical significance evident in the LS probe trials. This decrease in MI with more frequent stimulation suggests that inhibition immediately after stimulation was associated with firing rate fatigue.

### S11. ADDITIONAL ANALYSIS OF THE FIRING RATE FATIGUE

As described in Section II D of the main text, multiple animal trials with each probe type (LS: 7 animals, LD: 6 animals) were carried out to collect data to compare the degree of firing rate fatigue. Different power levels were applied to each emitter in each animal trial. To provide a fair comparison between the probes, we classified the animal trials into low and high power conditions for each probe and compared the induced inhibition strength (using the modulation index (MI) as a metric) after the 1 s CW stimulation between the power conditions. Table I summarizes the number of animal trials and the mean power with the power range used for each condition. The peak output intensity range at the specified output power was also noted for each power condition to show that the LD probes consistently exhibited higher peak output intensities compared to the LS probes, irrespective of the low or high power conditions. Despite this, the inhibition response is more statistically stronger for the LS probe as shown in Fig. 5. This supports our claim in the main text that the LS probes could induce stronger inhibition than the LD probes while operating at lower output intensities. The peak output intensity in Table I was estimated using the experimental approach described in Supplementary Section S5. The experimental approach was selected because it provided a more conservative intensity comparison between the two probe types, as evidenced by the smaller difference in peak output intensities between the LD and LS probes compared to the simulation approach (Simulation: 15.7*×*, Experiment: 9.7*×*).

**TABLE I.** Optical powers and intensities used in the animal trials.

| **Condition** | **Number of**<br>**Animals (n)** | **Average Power**<br>**($\mu\text{W}$)** | **Power Range**<br>**($\mu\text{W}$)** | **Intensity Range**<br>**($\text{mW}/\text{mm}^2$)** |
| --- | --- | --- | --- | --- |
| LD low power | 3 | 6.4 | 4.4 - 8.7 | 12.3 - 24.4 |
| LD high power | 3 | 10.5 | 6.7 - 13.9 | 18.7 - 39 |
| LS low power | 4 | 8.6 | 5.5 - 14.4 | 1.6 - 4.3 |
| LS high power | 3 | 21.5 | 14.7 - 26.7 | 4.4 - 8 |

**FIG. S12.**
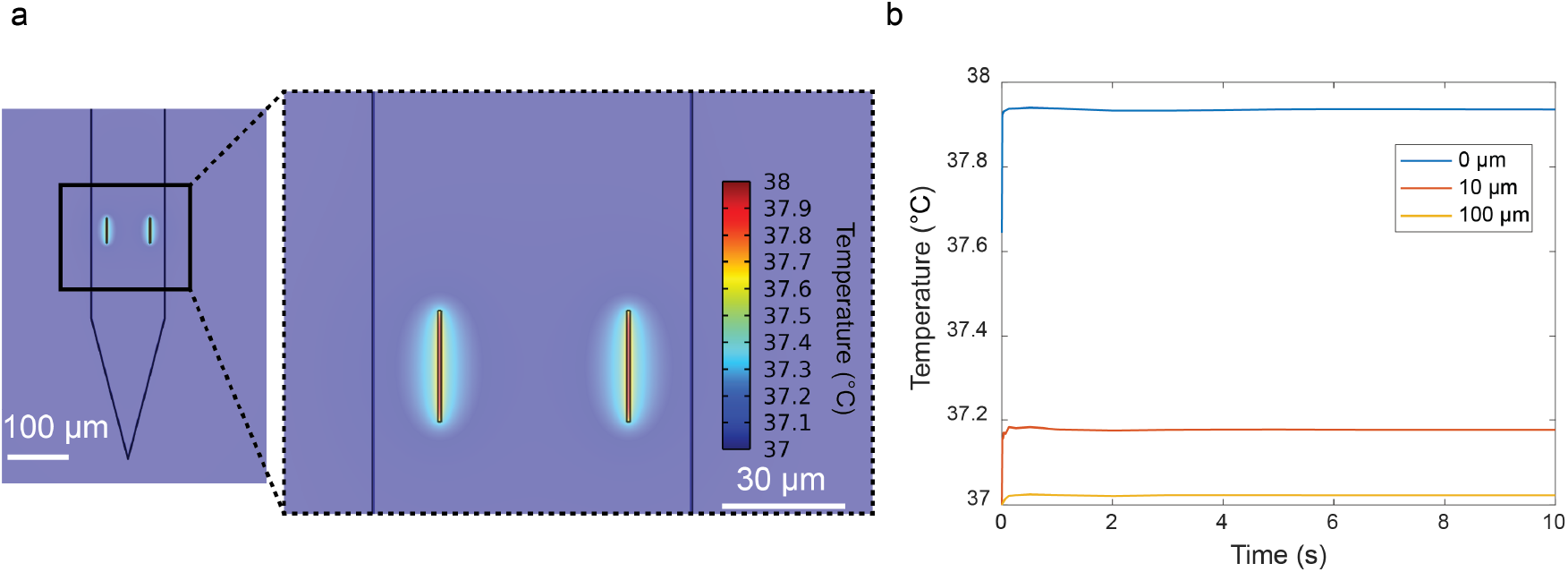
Simulation of the temperature increase in brain tissue due to optical absorption of 35 *µ*W from each grating emitter on an LS probe. **a** Simulated temperature distribution of the two emitters on a shank of an LS probe. The inset shows the zoom-in view of the same simulation result. **b** Change in temperature over 10 s CW output at 0, 10, and 100 *µ*m away from one of the emitters. The maximum temperature of the emitter reaches thermal equilibrium at 37.94 *^◦^C* in the brain.

Table II summarizes the pairwise statistical comparison of the MI. In general, a statistical difference was observed between the LS probe experiments and the LD low-power category, despite the higher peak output intensity from the LD probe. By increasing the output power of the LD probe from, on average, 6.4 *µ*W to 10.5 *µ*W, we noticed a decrease in the statistical significance of the MI difference between the LD and LS probe trials. This result indicates that the LD probe can potentially induce comparable inhibition responses as the LS probe at the cost of a higher peak output intensity. The average peak output intensity of the LD probe at the high power condition is *∼* 11*×* higher than the peak output intensity of the LS probe at the low power condition.

**TABLE II.**
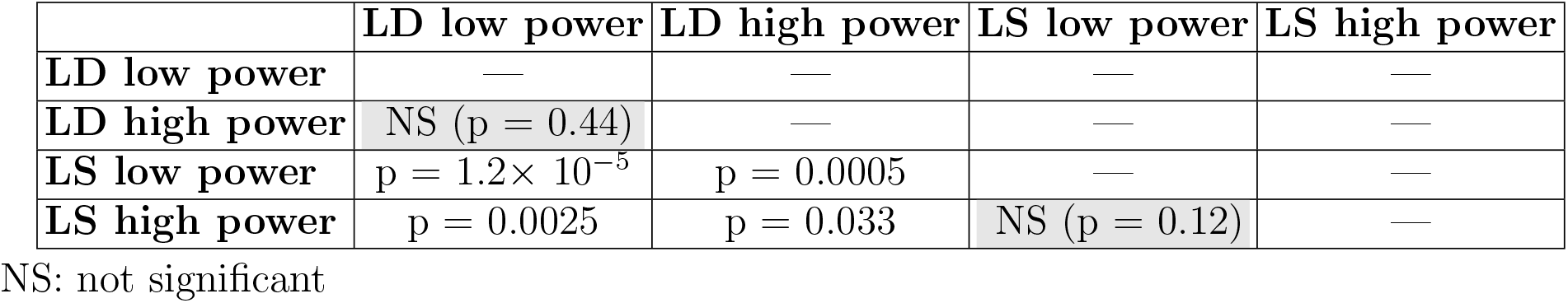
Statistical comparison (p-values) of the modulation index between 10 s before and 0.5 s after the 1 s CW stimulation using the two probe types at different output powers

|  | **LD low power** | **LD high power** | **LS low power** | **LS high power** |
| --- | --- | --- | --- | --- |
| **LD low power** | — | — | — | — |
| **LD high power** | NS (p = 0.44) | — | — | — |
| **LS low power** | p = $1.2 \times 10^{-5}$ | p = 0.0005 | — | — |
| **LS high power** | p = 0.0025 | p = 0.033 | NS (p = 0.12) | — |

NS: not significant

### S12. THERMAL SIMULATION OF THE OPTICAL ABSORPTION IN TISSUE

To investigate the optically induced temperature increase in brain tissue with the LS probe, we conducted thermal simulations using COMSOL Multiphysics. In this analysis, we determined the worst-case scenario for the temperature increase in the tissue by assuming that the emitted light is completely absorbed at the output of the light sheet emitter on the surface of the probe. We represented the heat generated by two grating emitters on each shank with two heat sources on the probe surface, each matching the size of the emitter. We set the power of each heat source to 35 *µ*W, equivalent to a total of 280 *µ*W of the entire light sheet. We employed the bioheat equation in COMSOL to model heat dissipation in the brain. The equation models the tissue as a solid medium with thermal conductivity *k_brain_*, density *ρ_brain_*, and heat capacity *C_p,brain_*. It also accounts for the cooling convection caused by blood perfusion and the heat generation from metabolism. Table III summarizes the parameters used in the simulation. These parameters are identical to those used in the thermal simulation to estimate the temperature increase with the µ-LED neural probe, as described in [S8].

**TABLE III.**
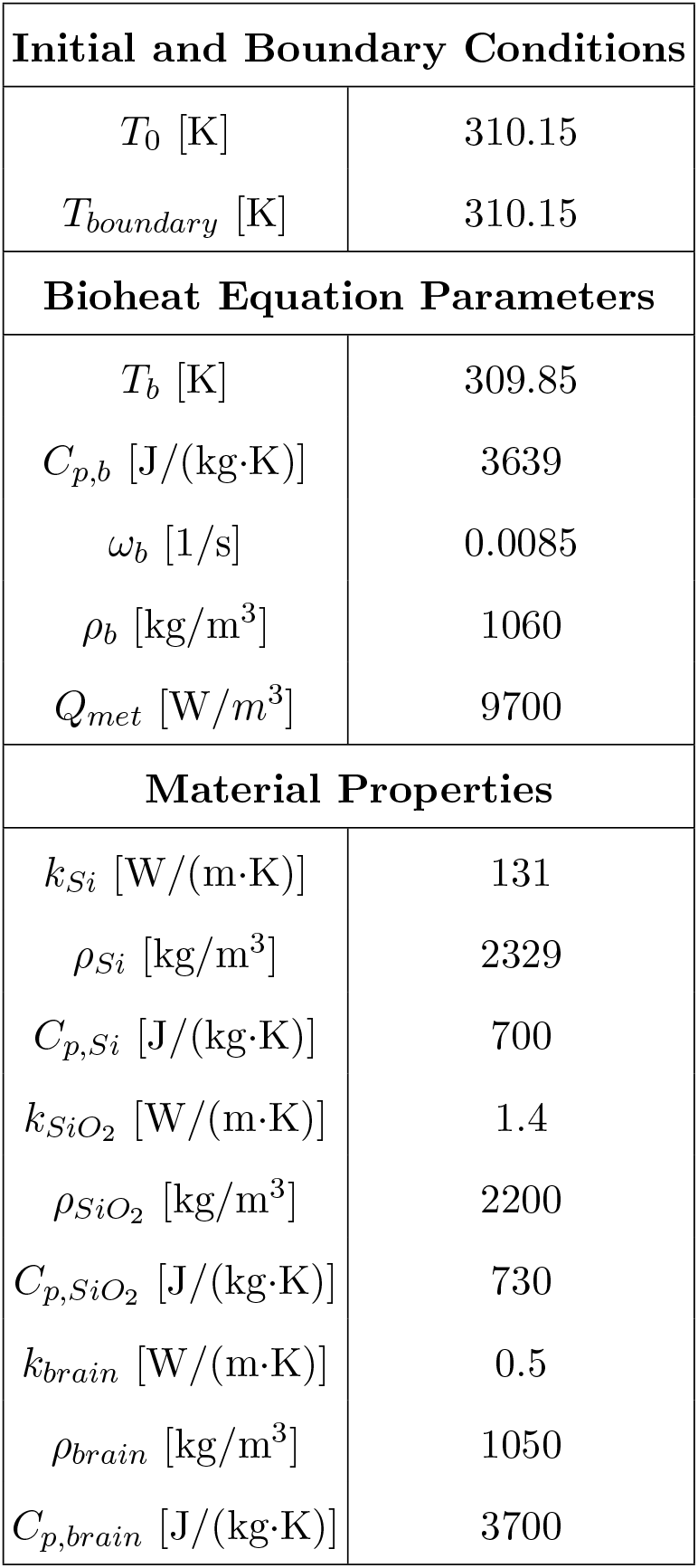
Summary of the parameters in the photothermal simulations.

| **Initial and Boundary Conditions** |  |
| --- | --- |
| $T_0$ [K] | 310.15 |
| $T_{boundary}$ [K] | 310.15 |
| **Bioheat Equation Parameters** |  |
| $T_b$ [K] | 309.85 |
| $C_{p,b}$ [J/(kg·K)] | 3639 |
| $\omega_b$ [1/s] | 0.0085 |
| $\rho_b$ [kg/m<sup>3</sup>] | 1060 |
| $Q_{met}$ [W/m<sup>3</sup>] | 9700 |
| **Material Properties** |  |
| $k_{Si}$ [W/(m·K)] | 131 |
| $\rho_{Si}$ [kg/m<sup>3</sup>] | 2329 |
| $C_{p,Si}$ [J/(kg·K)] | 700 |
| $k_{SiO_2}$ [W/(m·K)] | 1.4 |
| $\rho_{SiO_2}$ [kg/m<sup>3</sup>] | 2200 |
| $C_{p,SiO_2}$ [J/(kg·K)] | 730 |
| $k_{brain}$ [W/(m·K)] | 0.5 |
| $\rho_{brain}$ [kg/m<sup>3</sup>] | 1050 |
| $C_{p,brain}$ [J/(kg·K)] | 3700 |

Figure S12 illustrates that, at an output power of 35 *µ*W from each grating emitter, the temperature increase at the emitter surface remains within 1*^◦^*C. For this simulation, we considered only the thermal contribution from one of the four shanks, given that the increase in temperature at a distance of 100 *µ*m away from the probe is 0.02*^◦^C*. We expect the actual temperature increase of the brain at this output power to be lower than the simulated value because the optical absorption coefficient at a wavelength of 473 nm is *∼* 0.6 cm*^−^*^1^ [S9], so at least 1 mm of propagation distance is needed to absorb 20 % of the output power. The finite absorption leads to heat being generated over a larger area and a lower temperature increase in practice.

### S13. COMPARISON OF SI PHOTOSTIMULATION NEURAL PROBES

Table IV compares the probes in this work with state-of-the-art Si neural probes with photostimulation capabilities. In the table, wg stands for “waveguide.” The specifications are reported based on what has been demonstrated in *in vivo* experiments.

**TABLE IV.** Comparison of Si neural probes with photostimulation capability.

| Emitter type | No. of electrodes | No. of light emitters | Implant cross-section area [$\mu\text{m}^2$] | No. of shanks | Monolithically integrated light emitter? | Beam profile | Max. power output per emitter | Wave-length [nm] | Optical crosstalk | Foundry? | Ref. |
| --- | --- | --- | --- | --- | --- | --- | --- | --- | --- | --- | --- |
| SiN wg grating | 18 | 16 | $100 \times 58^a$ | 1 | Yes | Low-divergence | $\sim 200 \mu\text{W}$ | 488 | $< -35 \text{ dB}$<br>($\lambda = 488 \text{ nm}$) | Yes | This work (LD probe) |
| SiN wg grating | 56 | 5 | $78 \times 52^a$ | 4 | Yes | Planar sheet | $\sim 200 \mu\text{W}$ | 488 | $< -35 \text{ dB}$<br>($\lambda = 488 \text{ nm}$) | Yes | This work (LS probe) |
| SiN wg grating | 4 | 8 | $> 1000 \times 200$ | 1 | Yes | Low-divergence | Not reported | 473 | -13 dB | No | [S10] |
| SiN wg grating | 24 | 12 | $100 \times 30$ | 1 | Yes | Low-divergence | Not reported | 450 | None | Yes | [S11] |
| $\mu\text{-LED}$ | 256 | 128 | $144 \times 30$ | 4 | Yes | Lambertian | $\sim 7 \mu\text{W}$ | 470 | None | No | [S8] |
| $\mu\text{-OLED}$ | 32 | 1024 | $100 \times 55$ | 4 | No | Lambertian | $\sim 100 \text{ nW}$ | 490, 620<sup>b</sup> | None | No | [S12] |
| Multimode fiber | $960 \times 3$ probes | 1 | $70 \times 20 + 212.5^2 \times \pi^c$ | 1 | No | Diverging beam | $> 1 \text{ mW}$ | 450 - 495<sup>d</sup> | NA | Yes<sup>e</sup> | [S13, S14] |
| Polished fiber | 32 | up to 60 | 75 $\mu\text{m}$ wide probe<sup>f</sup> + $5.8^2 \times \pi$ per emitter<sup>c</sup> | 1 | No | Lambertian-like at fiber tip | $> 1 \text{ mW}$ | 473, 594 | $\sim -25 \text{ dB}$ | No | [S15] |

<sup>a</sup> Reporting the device with the smallest cross-section used in *in vivo* experiments.
<sup>b</sup> Peak emission wavelength estimated using Fig. 2d in [S12].
<sup>c</sup> The cross-section area of the electrophysiology silicon probe + the fiber.
<sup>d</sup> The wavelength used was not explicitly stated in the work. However, it was inferred to be in the blue spectrum since the optogenetic experiments were conducted with ChR2.
<sup>e</sup> Only the electrophysiology Si probe was fabricated in the foundry.
<sup>f</sup> Probe thickness not reported.

## References

[1] E. S. Boyden, F. Zhang, E. Bamberg, G. Nagel, and K. Deisseroth, Nature neuroscience 8, 1263 (2005).

[2] J. A. Cardin, M. Carlén, K. Meletis, U. Knoblich, F. Zhang, K. Deisseroth, L.-H. Tsai, and C. I. Moore, Nature protocols 5, 247 (2010).

[3] T.-W. Chen, T. J. Wardill, Y. Sun, S. R. Pulver, S. L. Renninger, A. Baohan, E. R. Schreiter, R. A. Kerr, M. B. Orger, V. Jayaraman, et al., Nature 499, 295 (2013).

[4] Y. Gong, C. Huang, J. Z. Li, B. F. Grewe, Y. Zhang, S. Eismann, and M. J. Schnitzer, Science 350, 1361 (2015).

[5] C. A. Baker, Y. M. Elyada, A. Parra, and M. M. Bolton, Elife 5, e14193 (2016).

[6] R. Lu, W. Sun, Y. Liang, A. Kerlin, J. Bierfeld, J. D. Seelig, D. E. Wilson, B. Scholl, B. Mohar, M. Tanimoto, et al., Nature neuroscience 20, 620 (2017).

[7] N. C. Pégard, A. R. Mardinly, I. A. Oldenburg, S. Sridharan, L. Waller, and H. Adesnik, Nature communications 8, 1228 (2017).

[8] W. Yang, L. Carrillo-Reid, Y. Bando, D. S. Peterka, and R. Yuste, elife 7, e32671 (2018).

[9] E. M. C. Hillman, V. Voleti, W. Li, and H. Yu, Annual review of neuroscience 42, 295 (2019).

[10] T. V. F. Abaya, S. Blair, P. Tathireddy, L. Rieth, and F. Solzbacher, Biomed. Opt. Express 3, 3087 (2012).

[11] M. M. Elwassif, Q. Kong, M. Vazquez, and M. Bikson, Journal of neural engineering 3, 306 (2006).

[12] J. M. Stujenske, T. Spellman, and J. A. Gordon, Cell reports 12, 525 (2015).

[13] S. F. Owen, M. H. Liu, and A. C. Kreitzer, Nature neuroscience 22, 1061 (2019).

[14] A. M. Aravanis, L.-P. Wang, F. Zhang, L. A. Meltzer, M. Z. Mogri, M. B. Schneider, and K. Deisseroth, Journal of neural engineering 4, S143 (2007).

[15] F. Pisanello, G. Mandelbaum, M. Pisanello, I. A. Oldenburg, L. Sileo, J. E. Markowitz, R. E. Peterson, A. D. Patria, T. M. Haynes, M. S. Emara, et al., Nature neuroscience 20, 1180 (2017).

[16] D. Eriksson, A. Schneider, A. Thirumalai, M. Alyahyay, B. de la Crompe, K. Sharma, P. Ruther, and I. Diester, Nature Communications 13, 985 (2022).

[17] A. M. Stamatakis, M. J. Schachter, S. Gulati, K. T. Zitelli, S. Malanowski, A. Tajik, C. Fritz, M. Trulson, and S. L. Otte, Frontiers in neuroscience, 496 (2018).

[18] N. Accanto, I.-W. Chen, E. Ronzitti, C. Molinier, C. Tourain, E. Papagiakoumou, and V. Emiliani, Scientific reports 9, 7603 (2019).

[19] J. H. Jennings, C. K. Kim, J. H. Marshel, M. Raffiee, L. Ye, S. Quirin, S. Pak, C. Ramakrishnan, and K. Deisseroth, Nature 565, 645 (2019).

[20] J. Zhang, R. N. Hughes, N. Kim, I. P. Fallon, K. Bakhurin, J. Kim, F. P. U. Severino, and H. H. Yin, Nature Biomedical Engineering 7, 499 (2023).

[21] F. Wu, E. Stark, P.-C. Ku, K. D. Wise, G. Buzsáki, and E. Yoon, Neuron 88, 1136 (2015).

[22] R. Scharf, T. Tsunematsu, N. McAlinden, M. D. Dawson, S. Sakata, and K. Mathieson, Scientific reports 6, 28381 (2016).

[23] K. Kim, M. Vöröslakos, J. P. Seymour, K. D. Wise, G. Buzsáki, and E. Yoon, Nature Communications 11 (2020), 10.1038/s41467-020-15769-w.

[24] M. Vöröslakos, K. Kim, N. Slager, E. Ko, S. Oh, S. S. Parizi, B. Hendrix, J. P. Seymour, K. D. Wise, G. Buzsáki, et al., Advanced Science 9, 2105414 (2022).

[25] A. J. Taal, I. Uguz, S. Hillebrandt, C.-K. Moon, V. Andino-Pavlovsky, J. Choi, C. Keum, K. Deisseroth, M. C. Gather, and K. L. Shepard, Nature Electronics, 1 (2023).

[26] E. Segev, J. Reimer, L. C. Moreaux, T. M. Fowler, D. Chi, W. D. Sacher, M. Lo, K. Deisseroth, A. S. Tolias, A. Faraon, and M. L. Roukes, Neurophotonics 4, 1 (2016).

[27] S. Libbrecht, L. Hoffman, M. Welkenhuysen, C. V. D. Haute, V. Baekelandt, D. Braeken, and S. Haesler, Journal of Neurophysiology 120, 149 (2018).

[28] A. Mohanty, Q. Li, M. A. Tadayon, S. P. Roberts, G. R. Bhatt, E. Shim, X. Ji, J. Cardenas, S. A. Miller, A. Kepecs, and M. Lipson, Nature Biomedical Engineering 4, 223 (2020).

[29] V. Lanzio, G. Telian, A. Koshelev, P. Micheletti, G. Presti, E. D’Arpa, P. D. Martino, M. Lorenzon, P. Denes, M. West, et al., Microsystems and Nanoengineering 7, 1 (2021).

[30] W. D. Sacher, F.-D. Chen, H. Moradi-Chameh, X. Luo, A. Fomenko, P. Shah, T. Lordello, X. Liu, I. F. Almog, J. N. Straguzzi, et al., Neurophotonics 8, 25003 (2021).

[31] W. D. Sacher, F.-D. Chen, H. Moradi-Chameh, X. Liu, I. F. Almog, T. Lordello, M. Chang, A. Naderian, T. M. Fowler, E. Segev, et al., Optics Letters 47, 1073 (2022).

[32] D. F. English, S. McKenzie, T. Evans, K. Kim, E. Yoon, and G. Buzsáki, Neuron 96, 505 (2017).

[33] M. Valero, I. Zutshi, E. Yoon, and G. Buzsáki, Science 375, 570 (2022).

[34] K. Kampasi, D. F. English, J. Seymour, E. Stark, S. McKenzie, M. Vöröslakos, G. Buzsáki, K. D. Wise, and E. Yoon, Microsystems and nanoengineering 4, 10 (2018).

[35] J. J. Jun, N. A. Steinmetz, J. H. Siegle, D. J. Denman, M. Bauza, B. Barbarits, A. K. Lee, C. A. Anastassiou, A. Andrei, Cağatay Aydn, et al., Nature 551, 232 (2017).

[36] L. Sileo, S. H. Bitzenhofer, B. Spagnolo, J. A. Pöpplau, T. Holzhammer, M. Pisanello, F. Pisano, E. Bellistri, E. Maglie, M. D. Vittorio, et al., Frontiers in neuroscience 12, 771 (2018).

[37] D. J. Calame, M. I. Becker, and A. L. Person, Nature Neuroscience, 1 (2023).

[38] E. G. McBride, S. R. Gandhi, J. R. Kuyat, D. R. Ollerenshaw, A. Arkhipov, C. Koch, and S. R. Olsen, Neuron 111, 275 (2023).

[39] T. il Kim, J. G. McCall, Y. H. Jung, X. Huang, E. R. Siuda, Y. Li, J. Song, Y. M. Song, H. A. Pao, R.-H. Kim, et al., Science 340, 211 (2013).

[40] W. D. Sacher, X. Luo, Y. Yang, F.-D. Chen, T. Lordello, J. C. C. Mak, X. Liu, T. Hu, T. Xue, P. G.-Q. Lo, M. L. Roukes, and J. K. S. Poon, Opt. Express 27, 37400 (2019).

[41] F.-D. Chen, Y. Jung, T. Xue, J. C. C. Mak, X. Luo, P. G.-Q. Lo, M. L. Roukes, J. K. S. Poon, and W. D. Sacher (2021) p. SW3B–2.

[42] A. Sharma, A. Govdeli, T. Xue, F.-D. Chen, X. Luo, H. Chua, G.-Q. Lo, W. D. Sacher, and J. K. S. Poon (2023) p. SF2E–5.

[43] T. Xue, A. Stalmashonak, P. Ding, W. D. Sacher, and J. K. S. Poon (2023) p. 1.

[44] S. S. Azadeh, A. Stalmashonak, K. W. Bennett, F.-D. Chen, W. D. Sacher, and J. K. S. Poon, Opt. Lett. 47, 26 (2022).

[45] J. P. Neto, P. Baião, G. Lopes, J. Frazão, J. Nogueira, E. Fortunato, P. Barquinha, and A. R. Kampff, Frontiers in neuroscience 12, 715 (2018).

[46] A. Berndt, P. Schoenenberger, J. Mattis, K. M. Tye, K. Deisseroth, P. Hegemann, and T. G. Oertner, Proceedings of the National Academy of Sciences 108, 7595 (2011).

[47] F. Fabbrini, C. den Haute, M. D. Vitis, V. Baekelandt, W. Vanduffel, and R. Vogels, Current Biology 29, 1988 (2019).

[48] K. Onodera and H. K. Kato, Nature Communications 13, 2585 (2022).

[49] Q. Wang, S. Yu, A. Simonyi, G. Y. Sun, and A. Y. Sun, Molecular neurobiology 31, 3 (2005).

[50] S. ichiro Osawa, M. Iwasaki, R. Hosaka, Y. Matsuzaka, H. Tomita, T. Ishizuka, E. Sugano, E. Okumura, H. Yawo, N. Nakasato, et al., PloS one 8, e60928 (2013).

[51] R. J. Racine, Electroencephalography and clinical neurophysiology 32, 281 (1972).

[52] B. Spagnolo, A. Balena, R. T. Peixoto, M. Pisanello, L. Sileo, M. Bianco, A. Rizzo, F. Pisano, A. Qualtieri, D. D. Lofrumento, F. D. Nuccio, J. A. Assad, B. L. Sabatini, M. D. Vittorio, and F. Pisanello, Nature Materials (2022), 10.1038/s41563-022-01272-8.

[53] G. Rios, E. V. Lubenov, D. Chi, M. L. Roukes, and A. G. Siapas, Nano letters 16, 6857 (2016).

[54] X. Cheng, Y. Li, J. Mertz, S. Sakadžić, A. Devor, D. A. Boas, and L. Tian, Optics letters 44, 4989 (2019).

[55] G. Yona, N. Meitav, I. Kahn, and S. Shoham, eneuro 3 (2016).

[56] A. Golabchi, “Wiring configuration chronic experiments,” (2021)

[57] R. Fiáth, A. L. Márton, F. Mátyás, D. Pinke, G. Márton, K. Tóth, and I. Ulbert, Scientific Reports 9, 111 (2019).

[58] Z. Zeidler, M. Brandt-Fontaine, C. Leintz, C. Krook-Magnuson, T. Netoff, and E. Krook-Magnuson, eneuro 5 (2018).

[59] P. Yger, G. L. B. Spampinato, E. Esposito, B. Lefebvre, S. Deny, C. Gardella, M. Stimberg, F. Jetter, G. Zeck, S. Picaud, et al., Elife 7, e34518 (2018).

[60] C. Rossant, A. Buccino, M. Economo, C. Gestes, D. Goodman, M. Hunter, S. Kadir, C. Nolan, M. Spacek, and N. Steinmetz, “phy: interactive visualization and manual spike sorting of large-scale ephys data,” (2020).

[61] N. Schmitzer-Torbert, J. Jackson, D. Henze, K. Harris, and A. D. Redish, Neuroscience 131, 1 (2005).

[62] S. Suner, M. R. Fellows, C. Vargas-Irwin, G. K. Nakata, and J. P. Donoghue, IEEE transactions on neural systems and rehabilitation engineering 13, 524 (2005).

[63] M. G. K. Brunk, K. E. Deane, M. Kisse, M. Deliano, S. Vieweg, F. W. Ohl, M. T. Lippert, and M. F. K. Happel, Scientific Reports 9, 20385 (2019).

[64] D. Yamamura, A. Sano, and T. Tateno, Brain Research 1659, 96 (2017).

[65] H. Shin, Y. Son, U. Chae, J. Kim, N. Choi, H. J. Lee, J. Woo, Y. Cho, S. H. Yang, C. J. Lee, et al., Nature communications 10, 3777 (2019).

## References

[S1] F.-D. Chen, H. Wahn, T. Xue, Y. Jung, J. N. Straguzzi, S. S. Azadeh, A. Stalmashonak, H. Chua, X. Luo, P. Shah, et al. (2022) p. JTh6A–7.

[S2] W. D. Sacher, X. Luo, Y. Yang, F.-D. Chen, T. Lordello, J. C. C. Mak, X. Liu, T. Hu, T. Xue, P. G.-Q. Lo, M. L. Roukes, and J. K. S. Poon, Opt. Express 27, 37400 (2019).

[S3] W. D. Sacher, F.-D. Chen, H. Moradi-Chameh, X. Liu, I. F. Almog, T. Lordello, M. Chang, A. Naderian, T. M. Fowler, E. Segev, et al., Optics Letters 47, 1073 (2022).

[S4] W. D. Sacher, F.-D. Chen, H. Moradi-Chameh, X. Luo, A. Fomenko, P. Shah, T. Lordello, X. Liu, I. F. Almog, J. N. Straguzzi, et al., Neurophotonics 8, 25003 (2021).

[S5] Y. Lin, J. C. C. Mak, H. Chen, X. Mu, A. Stalmashonak, Y. Jung, X. Luo, P. G.-Q. Lo, W. D. Sacher, and J. K. S. Poon, Optics Express 29, 34565 (2021).

[S6] X. Mu, F.-D. Chen, K. M. Dang, M. G. K. Brunk, J. Li, H. Wahn, A. Stalmashonak, P. Ding, X. Luo, H. Chua, et al., Front. Neurosci. 17, 1213265 (2023).

[S7] M. Gaidica, “Mouse brain atlas,” (2001), accessed: 2023-09-20.

[S8] M. Vöröslakos, K. Kim, N. Slager, E. Ko, S. Oh, S. S. Parizi, B. Hendrix, J. P. Seymour, K. D. Wise, G. Buzsáki, et al., Advanced Science 9, 2105414 (2022).

[S9] G. Yona, N. Meitav, I. Kahn, and S. Shoham, eneuro 3 (2016).

[S10] A. Mohanty, Q. Li, M. A. Tadayon, S. P. Roberts, G. R. Bhatt, E. Shim, X. Ji, J. Cardenas, S. A. Miller, A. Kepecs, and M. Lipson, Nature Biomedical Engineering 4, 223 (2020).

[S11] S. Libbrecht, L. Hoffman, M. Welkenhuysen, C. V. D. Haute, V. Baekelandt, D. Braeken, and S. Haesler, Journal of Neurophysiology 120, 149 (2018).

[S12] A. J. Taal, I. Uguz, S. Hillebrandt, C.-K. Moon, V. Andino-Pavlovsky, J. Choi, C. Keum, K. Deisseroth, M. C. Gather, and K. L. Shepard, Nature Electronics, 1 (2023).

[S13] E. G. McBride, S. R. Gandhi, J. R. Kuyat, D. R. Ollerenshaw, A. Arkhipov, C. Koch, and S. R. Olsen, Neuron 111, 275 (2023).

[S14] J. J. Jun, N. A. Steinmetz, J. H. Siegle, D. J. Denman, M. Bauza, B. Barbarits, A. K. Lee, C. A. Anastassiou, A. Andrei, Caǧatay Aydn, et al., Nature 551, 232 (2017).

[S15] D. Eriksson, A. Schneider, A. Thirumalai, M. Alyahyay, B. de la Crompe, K. Sharma, P. Ruther, and I. Diester, Nature Communications 13, 985 (2022).

